# Reconciling conflicting selection pressures in the plant collaborative non-self recognition self-incompatibility system

**DOI:** 10.1101/2024.11.11.622984

**Authors:** Amit Jangid, Keren Erez, Ohad Noy Feldheim, Tamar Friedlander

**Affiliations:** The Robert H. Smith Institute of Plant Sciences and Genetics in Agriculture, Faculty of Agriculture, food and environment, The Hebrew University of Jerusalem, Rehovot, Israel; Einstein Institute for Mathematics, The Hebrew University of Jerusalem, Jerusalem, Israel

## Abstract

Complex biological systems should often reconcile conflicting selection pressures. In systems based on molecular recognition, molecules must specifically identify certain partners while excluding others. Here we study how such selection pressures shape the evolution of the self-incompatibility system in plants. This system inhibits self-fertilization using specific molecular recognition between proteins, expressed in the plant female and male reproductive organs. We study the impact of these opposing selection pressures on the amino acid frequencies in these proteins’ recognition domain. We construct a theoretical framework enabling promiscuous recognition between proteins, as found empirically, and employ stochastic simulations to study its evolution. We find that selection exerts asymmetric responses of amino acid frequencies, affecting female proteins considerably, but hardly the male. Using large deviations theory, we well-approximate the simulated frequencies and find agreement with genomic data. Our work offers a general theoretical framework to study the impact of multiple selection pressures, applicable to additional biological systems.

## Introduction

Multi-component biological systems frequently evolve under the influence of multiple, often conflicting, selection pressures acting on their components. These pressures shape the interactions between different components and ultimately contribute to the system’s overall function and survival. One example of such a system is the self-incompatibility mechanism in plants, which evolved to prevent self-fertilization and promote outcrossing to enhance genetic diversity [1, 2, 3, 4, 5, 6, 7, 8, 9]. Under this mechanism, the plant population is sub-divided into a few dozen ‘types’ or ‘classes’, such that a pollen grain cannot fertilize a maternal plant of the same type, but only plants of different types. The type is encoded by several ‘type-specifying’ genes, located in a single highly diverse genomic region called ‘S-haplotype’, and expressed in either the male or the female reproductive organs of the plant. These type-specifying genes are an excellent example of a medium-sized biological system, whose components should co-evolve in a concerted fashion to jointly achieve a unified functionality. Since this system’s functionality is tightly coupled to plant reproduction, the selection pressures acting on its components are strong. These considerations render the self-incompatibility system an excellent case study to explore how complex biological systems evolve under the constraints of competing selection pressures.

Self-incompatibility requires molecular recognition that could distinguish between different molecular types [10]. We concentrate on a particular self-incompatibility mechanism, relying on non-self-recognition. Namely, fertilization is disabled by default and enabled only if the incoming pollen is positively identified as having a non-self type. Thus, non-self-recognition requires each pollen to be equipped with multiple molecular ‘identifiers’, to allow fertilization of multiple non-self female types (Fig. 1a).

**Figure 1.**
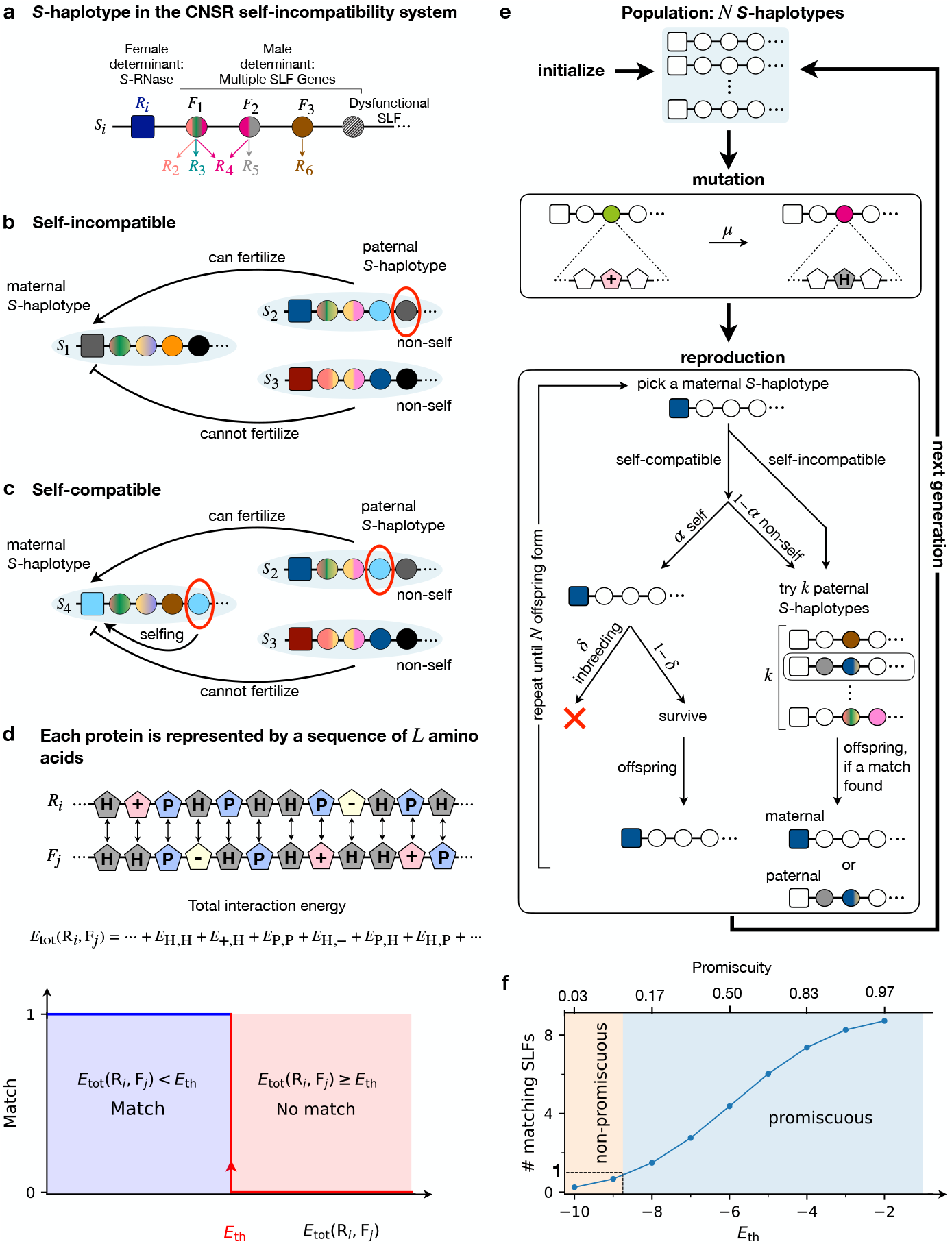
Model description. **(a)** Each S-haplotype consists of a single female type-specifying gene (RNase *R*_*i*_, square) and multiple male-type-specifying genes (SLFs *F*_1_, *F*_2_, …, circles). Each SLF could match one or several different RNases or none. **(b)** For a paternal S-haplotype to successfully fertilize a maternal S-haplotype it must be equipped with an SLF matching the maternal RNase. A maternal S-haplotype can be fertilized by either self (genetically identical) or non-self pollen. An S-haplotype is considered ‘self-incompatible’ if it does not contain an SLF matching its own RNase, rendering its fertilization by self-pollen impossible, and ‘self-compatible’ **(c)** if it does contain such an SLF facilitating fertilization by self-pollen. **(d)** Each protein encoded in the S-haplotype is represented by a sequence of *L* amino acids of four biochemical classes. The total interaction energy 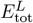 between an RNase *R*_*i*_ and each SLF *F*_*j*_ is defined as the sum of the pairwise interaction energies between their corresponding amino acids. Two such proteins are considered matching, if 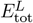 is smaller than an energy threshold *E*_th_, and non-matching otherwise. **(e)** The population life cycle implemented in our stochastic simulation. The population consists of a fixed number *N* of S-haplotypes, as shown in (a). Every generation, each of the amino acids in each of the proteins can mutate with probability *µ*. We assume non-overlapping generations. In every generation, maternal S-haplotypes are picked at random for reproduction. If the maternal S-haplotype chosen is self-compatible it can be fertilized by either non-self-pollen (with probability 1 − *α*) or by self-pollen (with probability *α*), where self-fertilization offspring survive with probability 1 − *δ* relative to offspring formed by non-self fertilization. Alternatively, if the maternal plant is self-incompatible only non-self pollination is possible. Under non-self pollination, the maternal plant is given up to *k* attempts to find a compatible paternal S-haplotype, by randomly picking S-haplotypes from the entire population. If a match is found, an offspring inheriting either the paternal or the maternal S-haplotype is generated. **(f)** We consider ‘promiscuity’ as the state that on average more than one SLF per S-haplotypes matches an arbitrary RNase. Here we plot the mean number of matching SLFs (assuming 9 SLFs per S-haplotype, our default value), against *E*_th_, where all sequences are randomly drawn from the prior distribution. We find that the system is promiscuous for *E*_th_ ≥ −9 (see also Table 3).

The female-side protein in this self-incompatibility mechanism, expressed in the female organs, is an RNase, where different types have different RNase alleles [11]. The RNase is toxic to the incoming pollen and by default inhibits fertilization. The male-side proteins, expressed in the pollen, are F-box proteins (called SLF), which are involved in targeted protein degradation [12, 13]. Each S-haplotype encodes multiple different SLF genes. To enable fertilization, at least one of the SLFs in the incoming pollen, should specifically match the female’s RNase molecules and mediate their degradation, allowing the pollen to evade the RNase harmful effect. If the RNase is successfully degraded and fertilization ensues, we say that the paternal S-haplotype (pollen) is compatible with the maternal S-haplotype (pollen recipient). Otherwise, these S-haplotypes are considered incompatible. Multiple SLF genes are encoded in the S-haplotype to collaboratively allow fertilization of multiple non-self types, granting this self-incompatibility mechanism the name the ‘collaborative non-self recognition’ (CNSR) self-incompatibility mechanism [14].

To avoid self-fertilization, an S-haplotype must not contain an SLF matching its own (self) RNase (Fig. 1b). A plant carrying an S-haplotype that includes such an SLF is capable of self-fertilization and considered ‘self-compatible’ (Fig. 1c). To avoid an accidental insertion of an SLF gene that could cause self-compatibility, and preserve an SLF combination capable of fertilizing multiple female types, recombination in this genomic region is nearly suppressed. As a result, the entire S-haplotype functions as a heritable evolutionary unit, balancing the selective pressures acting on its constituent genes.

Detailed mapping of the molecular recognition between various protein pairs uncovered that an SLF protein could match a variable number of distinct RNases [14, 15], but how exactly the specificities of these proteins are determined remains elusive. Computational studies estimated that only 10-20 residues determine the RNase [16, 17] and SLF specificities [18, 19, 20], and in one case the substitution of only three residues in an RNase was sufficient to modify its compatibility phenotype [21, 22]. Given that these protein specificities are governed by very few amino acids, it remains unclear how they reconcile the multiple and possibly conflicting selection pressures, to avoid self-compatibility and enhance cross-compatibility. Addressing this question requires an explicit model of the molecular recognition implemented by these proteins. Yet, previous studies [23, 24] employed simplified descriptions of these proteins, that did not explicitly include their interaction domain, but only attributed each protein a particular compatibility phenotype. As these models lack a description of the biophysical basis of the match between the proteins, they are unsuitable for studying the selection pressures shaping these proteins’ compatibility phenotypes. Two-tier models describing both the genotype and the phenotype levels of a biological system were previously constructed to describe RNA secondary structures [25, 26, 27], the immune system and olfactory receptors [28] and gene regulatory networks [29, 30]. All these cases share the notion of a many-to-one mapping between genotype to phenotype, proposed to be a general guiding principle of biological systems [31]. Underlying this degenerate mapping is the existence of numerous hidden microscopic degrees of freedom of the system. Evolutionary constraints operating at the phenotype level can often be satisfied by a variety of distinct genotypes, making the study of these internal degrees of freedom far more informative than focusing on the phenotype alone. For example, analyses of selection pressures shaping the spectrum of immune receptor repertoire [32] used detailed descriptions of their genotypes and evolutionary trajectories.

Inspired by these ideas, we have recently proposed a novel theoretical framework of the non-self recognition self-incompatibility system, that incorporates the molecular recognition between the male and female type-specifying proteins into the evolutionary model to study the emergence and maintenance of mating specificities [33] (see Fig. 1). The model uniquely distinguishes between the genotype and phenotype levels of the system and incorporates degenerate mapping from genotype to phenotype. Notably, our approach does not require the definition of arbitrary notions of fitness but rather derives fitness from the compatibility phenotype which directly depends on the interaction energies between the male and female type-specifying proteins. This modeling framework crucially relies on interaction promiscuity and facilitates many-to-many interactions between the female and male type-specifying proteins, in agreement with empirical evidence [14, 15]. Using this framework, we found that most S-haplotypes in our model spontaneously self-organize into ‘compatibility classes’, such that members of each class are incompatible with each other but compatible with all members of all other classes, as found empirically. We demonstrated a dynamic balance between class birth and death in this model, reaching a dynamically stable equilibrium in the class number, and researched the mutational trajectories by which class birth and death occur, facilitating the emergence and decay of mating specificities. The facilitation of neutral mutations in our model exposed new evolutionary trajectories for class emergence that could not be envisioned in simpler models. These desired properties of the model and, in particular, the explicit inclusion of molecular recognition into an evolutionary model render it highly suitable to study how the selection pressures shape these proteins’ specificities.

Here we employ this model to research the balance between the multiple selection pressures acting on the male and female type-specifying proteins in this system, as reflected by the amino acid frequencies in their specificity-determining residues. We analyze the amino acid frequencies of single amino acids in the RNase and SLF separately and the joint frequencies of corresponding amino acid pairs in both. Using Sanov’s Theorem from large deviations theory, we derive the amino acid frequencies as the maximizers of the entropy under a self incompatibility constraint. We demonstrate that a reduced model subject only to selection against self-compatibility closely approximates the frequencies obtained in the full model. This result highlights the dominance of selection against self-compatibility in the evolution of this system. To disentangle the impacts of attractive vs. repulsive selection pressures we distinguish between same-S-haplotype and cross-S-haplotype gene pairs. We identify two strategies employed by these sequences in response to the combination of both selection pressures: a ‘global’ strategy – where proteins are broadly attractive or repulsive regardless of the specific partner, and a ‘local’ strategy – where proteins selectively match or mismatch only particular partners. A ‘global’ strategy is implemented by biases in single amino acid frequencies, whereas a ‘local’ strategy relies on specific geometric adjustment between particular sequence pairs, reflected in biases in their joint amino acid frequencies. We show that avoidance of self-compatibility is mostly achieved by a global strategy via decrease in hydrophobic and increase in charged amino acid frequencies in the female type-specifying proteins. In contrast, compatibility with partners is often obtained via a local strategy of matching the locations of positively and negatively charged amino acids. Lastly, we validate our theoretical

## Results

### The model

We assume a population of *N* S-haplotypes, each consisting of a single RNase and multiple SLF genes (Fig. 1a). Each of the proteins encoded by these genes is represented by a sequence of *L* amino acids, representing the protein binding interface, that could belong to one of four biochemical classes: hydrophobic (H), neutral polar (P), positively charged (+), or negatively charged (–). An RNase-SLF pair is considered matching if the interaction energy between their interfaces is below a threshold value *E*_th_. This interaction energy between an RNase *R*_*i*_ and SLF *F*_*j*_ is defined as the sum of the pairwise interaction energies between the corresponding amino acids (see Table 1) of these proteins,

**Table 1:** Interaction energies between pairs of amino acids.

|  | H | P | + | - |
| --- | --- | --- | --- | --- |
| Hydrophobic (H) | -1 | 0 | 0 | 0 |
| Neutrally polarized (P) | 0 | -0.75 | -0.25 | -0.25 |
| Positively charged (+) | 0 | -0.25 | 1 | -1.25 |
| Negatively charged (-) | 0 | -0.25 | -1.25 | 1 |

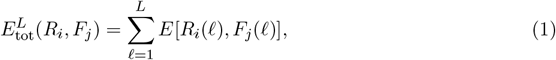

where *R*_*i*_(*l;*), *F*_*j*_(*l*) ∈ {H, P, +, −}. If 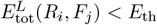, these proteins are considered matching, namely fertilization of a maternal S-haplotype carrying the RNase *R*_*i*_ by a paternal S-haplotype carrying the SLF *F*_*j*_ is enabled (Fig. 1d). Hence, the S-haplotype composition determines its possible mating partners. In particular, an S-haplotype is termed ‘self-incompatible’ if it does not contain any SLF that matches its own RNase (Fig. 1b) and ‘self-compatible’ if it does (Fig. 1c). Self-compatible S-haplotypes can be fertilized by either self or non-self pollen, whereas self-incompatible ones can only be fertilized by non-self pollen.

Each individual produces both male and female gametes, but the male gametes typically outnumber the female ones. This natural asymmetry in abundance is captured in our model by the number of fertilization attempts *k >* 1 per female. We assume a well-mixed population with no spatial structure, such that every female is equally likely to encounter pollen from any other population member. Every female gamete receives non-self-pollen originating uniformly from any other population member, as well as self-pollen coming from the same plant. The results below refer to the regime of pollen abundance *k* ≫ 1 in which a female gamete is essentially guaranteed to be fertilized, and only the male gametes compete for matching female ones.

The population life-cycle is illustrated in Fig. 1e. We initialize a population of *N* S-haplotypes, constructed as shown in Fig. 1a,d. We assume non-overlapping generations. Every generation, each of the amino acids in each of the proteins can be mutated with some probability. The new values of amino acids chosen to be mutated are randomly drawn from the prior amino acid distribution (Methods). In each generation, the offspring population is produced one by one from the parental population by repeating the following procedure. An S-haplotype is selected uniformly at random from the parental population to serve as a maternal parent. If the maternal S-haplotype is self-compatible, it is self-fertilized with probability *α* (here *α* corresponds to the proportion of self-pollen out of the total pollen received). If self-fertilization did not occur (whether or not the S-haplotype is self-compatible), the maternal S-haplotype is given *k* opportunities for fertilization by cross-pollination. In each of these fertilization opportinities, a random S-haplotype of the parental population is selected and its compatibility as male with the maternal parent is tested. If compatible, it serves as a paternal parent, and an offspring S-haplotype is formed. This offspring is an identical copy of either of its two parents, with equal probability. To represent the deficiency of offspring produced via self-fertilization, such offspring are eliminated with probability *δ* once produced. The parameter *δ* represents the inbreeding depression intensity. Hence every such round can either result in a single offspring, or none (due to failure of all fertilization attempts, or elimination of an offspring produced via self-fertilization). This procedure is repeated until a population of *N* offspring is formed. The offspring population then replaces the parental one, completing one generation.

Following this life-cycle, the population evolves under two types of selection pressures: avoidance of self-compatibility exerted by the inbreeding depression penalty on offspring produced via self-fertilization and selection for reproductive success where S-haplotypes compete on being compatible with as many non-self population members as possible. This model is a haploid version of the diploid model presented in [33]. Although simplified, the haploid model behaves qualitatively very similarly to the more realistic diploid model (see detailed comparison in supplementary text), and we opt for it due to its analytical and numerical tractability.

### The interaction energies between protein pairs vary with *E*_th_

A fundamental parameter of our model is *E*_th_– the energy threshold below which sequences are considered matching. *E*_th_ can alternatively be interpreted as representing the probability that protein pairs drawn at random from the prior distribution match each other. In the following we test the model behavior under a broad range of *E*_th_ values, spanning match probabilities between 0.03 to 0.97. Intuitively, we consider the population state as ‘promiscuous’, if on average an S-haplotype drawn at random from the prior distribution, contains more than a single SLF matching an arbitrary RNase (see Fig. 1f and Table 3).

The RNase and SLF sequences in our model evolve under a combination of two types of selection pressures, acting simultaneously to avoid match between RNase and SLF within the same S-haplotype but promote a match between RNase and SLF pairs located on different S-haplotypes – see Fig. 2 for schematic description.

**Figure 2:**
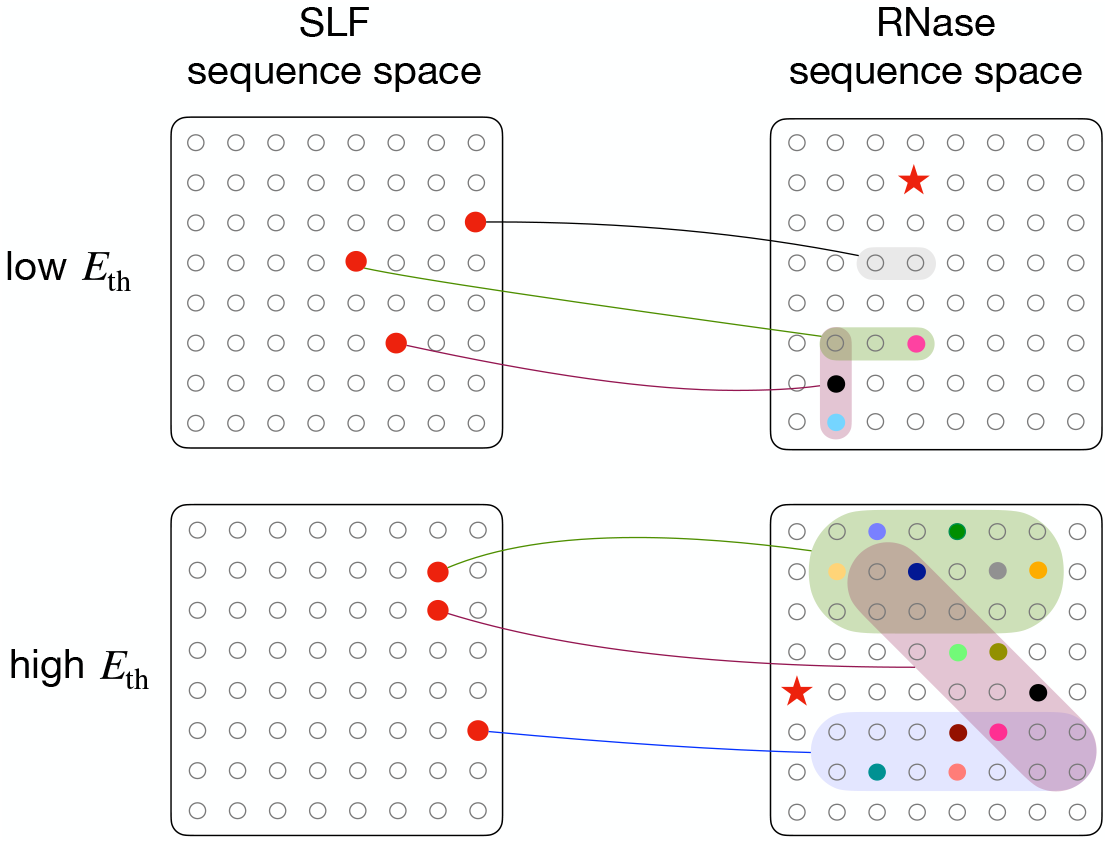
Match and mismatch relations between RNase and SLF proteins in a population of self-incompatible individuals – Schematic description. We use an abstract illustration of the high dimensional sequence space of the proteins encoded in the S-haplotype, where each circle represents a potential sequence: SLFs on the left and RNases on the right. Every SLF in a particular S-haplotype (red circle in the left panels) matches various RNase sequences (shaded region in the right panel). The self-RNase is shown as a red star and all the non-self RNases currently existing in the population are shown as circles in different colors. Collectively, these SLFs should match all the non-self RNases, but avoid matching the self-RNase. Empty circles represent RNase sequences not existing in the current population (right side) or SLF sequences not existing in the S-haplotype under study (left side). The upper part illustrates low promiscuity, where every SLF matches only a few RNase sequences, and the RNase diversity is low. The lower part depicts high promiscuity, where every SLF matches many RNases and the total number of distinct RNases in the population is higher. It is also possible that an SLF matches none of the current population RNases (shaded region with no colored circle), as appears in the upper part. Such an SLF is considered dysfunctional.

Since matches between proteins are determined by the interaction energies, the entire population sequences should co-evolve to collectively satisfy these match and mismatch constraints.

In Fig. 3a we present the interaction energies between RNase-SLF pairs obtained in simulation of the model for three different *E*_th_ values. We illustrate two types of RNase-SLF pairs exhibiting distinct interaction phenotypes: pairs encoded on the same S-haplotype (‘self’, red) that evolved to mismatch, and partners, i.e matching RNase and SLF pairs, such that the paternal S-haplotype carrying the SLF can fertilize the maternal S-haplotype carrying the RNase (‘partner’, blue). In all cases, the ‘self’ energy distribution is fully above *E*_th_, whereas the ‘partner’ distribution is entirely below *E*_th_, which sets the boundary between match and mismatch. We also show for reference the interaction energy distribution for sequences chosen at random from the prior distribution (‘neutral’, gray). Following the change in *E*_th_ we observe a shift in the interaction energy distributions relative to the neutral case, suggesting a role for the selection pressures acting on these sequences. The relative parts of the opposing selection pressures acting to attract and to repel should also depend on *E*_th_. Since the likelihood of a coincidental match increases with *E*_th_, the selection pressure to repel increases with *E*_th_ and the selection pressure to attract decreases. An additional feature of our model is the ability of each sequence to match multiple distinct partners. This number of partners also grows with *E*_th_. In Fig. 3b we show the distribution of the number of RNase partners per SLF as obtained in simulation for several *E*_th_ values. We observe a shift in the partner number distribution with *E*_th_ value, such that for high promiscuity the mean number of partners is larger and the distribution is broader, whereas for low promiscuity, the mean number of partners is smaller. Interestingly, for intermediate-to-low promiscuity, (*E*_th_ ≤ −6) a significant proportion of SLFs have no partner at all. Such dysfunctional SLFs likely survive by hitchhiking to linked functional ones.

**Figure 3.**
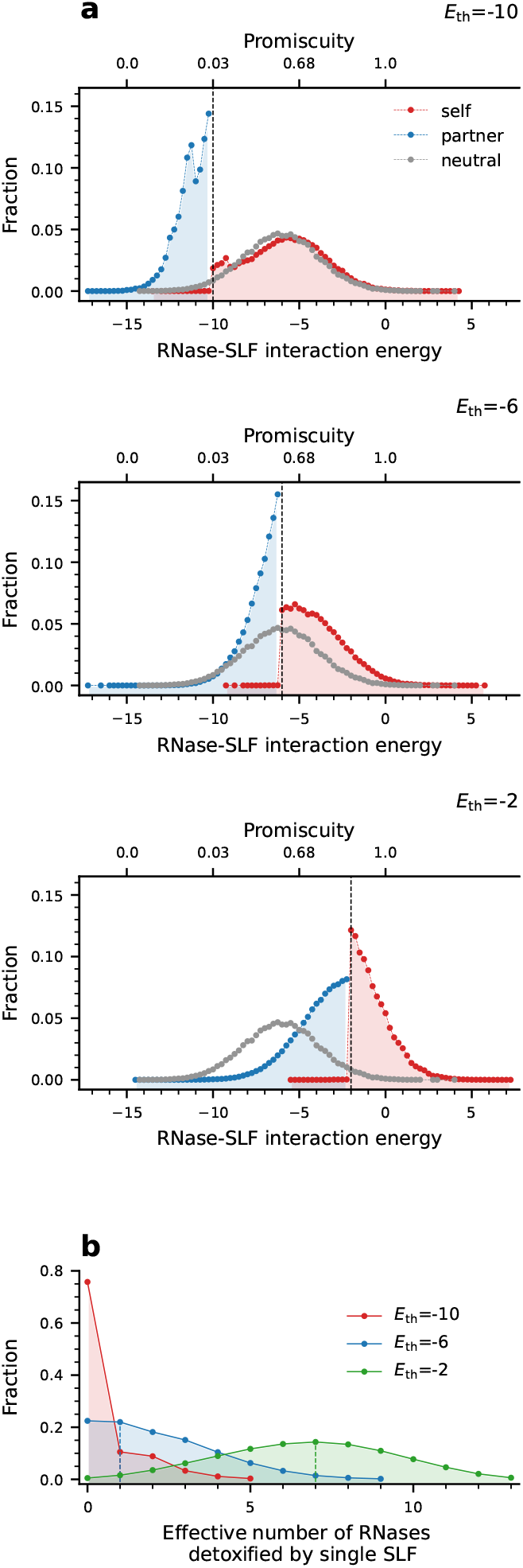
The typical interaction energies and the number of partners per protein vary with the interaction energy threshold *E*_th_. This figure shows steady-state characteristics of the population following evolution via our model. **(a)** Distributions of RNase-SLF interaction energies at steady state amongst populations evolved under three different *E*_th_ values. We illustrate separately interactions between two types of RNase-SLF pairs: same-haplotype RNase and SLF (‘self’, red); and interactions between partners - RNase-SLF pairs matching each other (and essentially located on S-haplotypes affiliated with different mating specificities) (‘partner’, blue). ‘Self’ pairs evolve to avoid match, hence their interaction energies exceed *E*_th_, and ‘partner’ pairs evolve to match, hence their energies are below the threshold. For reference, we also show the neutral distribution of RNase-SLF interaction energies, obtained for sequences drawn at random from the prior distribution (grey). *E*_th_ values are marked by a vertical dashed line. **(b)** Distributions of the effective number of distinct RNases in the population that a single SLF matches for different *E*_th_ values. These results refer to RNases and SLFs co-evolving in the same population. We observe that the distribution broadens and its mean increases with *E*_th_. SLFs with zero partners are considered dysfunctional, and their proportion increases as *E*_th_ decreases.

### Avoidance of self-compatibility governs the amino acid composition of the RNases, while the SLF composition remains nearly neutral

In the following section, we investigate the molecular underpinnings of the shift in interaction energies propelled by selection (Fig. 3). Below we show that the dominant selection pressure shaping the amino acid frequencies is avoidance of self-compatibility. To show that, we analyze the single amino acid frequencies in the evolved RNases and SLFs compared to neutral ones. To quantify the impact of selection relative to neutrality, we employ an information-theoretic measure of distance between distributions to compare the amino acid frequencies obtained under selection and in its absence. We then demonstrate that a reduced model with only selection against self-compatibility, but no selection for cross-compatibility, provides a good approximation to the AA frequencies in the full model.

We begin by examining the match probabilities between evolved RNase-SLF pairs, for different *E*_th_ values, compared to neutral sequences. The evolved sequences were obtained in simulation following the life cycle of Fig. 1 and were extracted after an initial transient time was discarded from the analysis, such that the population state is essentially independent of its initial condition and its statistical measures have already stabilized (see Fig. S1). We refer below to the distributions obtained as being at steady-state (see Methods). We find (Fig. 4a) that the evolved sequences (red) behave very differently from the neutral ones (blue), and for most of the *E*_th_ range are less likely to match relative to the neutral case. We conclude that for a great part of *E*_th_ values, a match at random is highly expected, and on average the selection pressure operates to reduce it.

**Figure 4:**
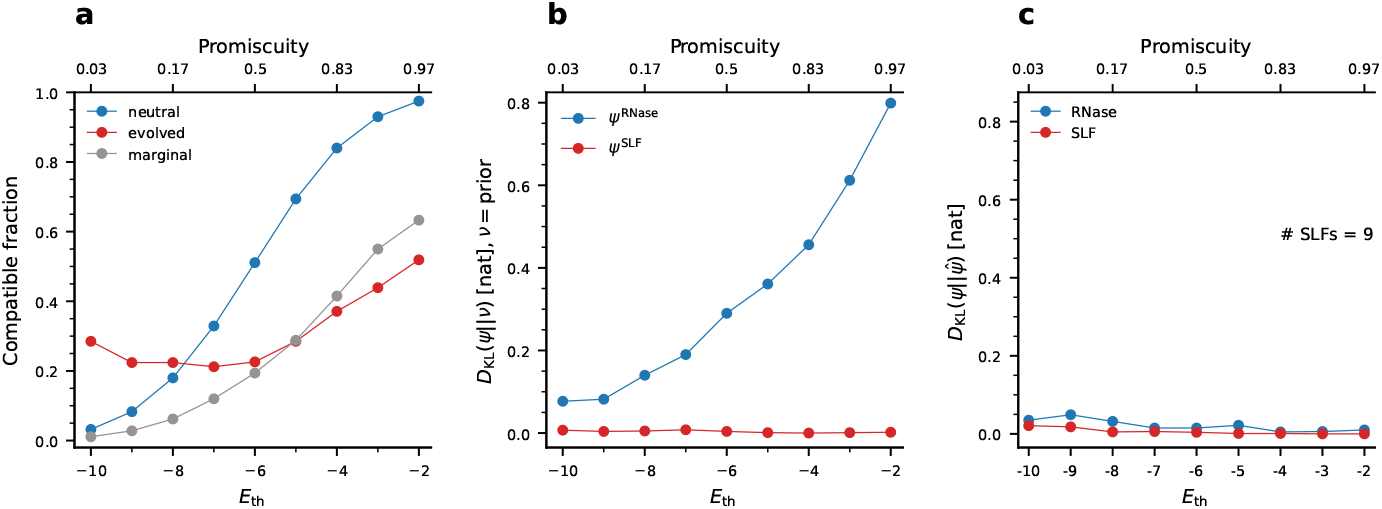
Avoidance of self-compatibility, biases the amino acid composition for RNases, but not for SLFs. **(a)** Compatible fraction amongst RNase-SLF pairs against *E*_th_ under three scenarios: neutral sequences drawn from the prior AA frequencies (blue), functional RNase and SLF sequences evolved in simulation (red), and random sequences drawn from the marginal AA frequencies of the evolved RNases and functional SLFs (gray). To calculate the neutral and marginal frequency cases, we drew a total of 50 RNases, and for each, we drew 1500 SLFs. **(b)** The Kullback-Leibler divergence between the marginal AA distribution of evolved sequences and the prior AA distribution for RNases (blue) and functional SLFs (red) as a function of *E*_th_. The calculations are based on 25 independent runs, using 600 data points from each run with 25 generation intervals between consecutive time points (Methods). **(c)** The divergence between the marginal distributions evolved under the full (*ψ*) and reduced 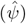 models, is close to zero, demonstrating that the reduced model provides an excellent approximation of the AA frequencies in the entire range of *E*_th_. Parameter values are as is Table 2.

We denote the prior distribution from which neutral sequences are drawn by *ν*(*i*) for *i* ∈ {+, −, *H, P*}, the marginal distributions of single amino acid frequencies in the evolved sequences by *ψ*^RNase^(*i*), *ψ*^SLF^(*i*) respectively, and the joint distribution of amino acid pairs in the same sequence position in RNase and SLF located on the same-S-haplotype by *φ*(*i, k*) for *i, k* ∈ {+, −, *H, P*}.

**Table 2:** Description of parameters and default values in the main model simulation.

| Parameters | Descriptions |
| --- | --- |
| $N = 2000$ | Total number of S-haplotypes or population size |
| $n = 10$ | Initial number of distinct S-haplotypes |
| $m = 9$ | Number of SLF paralogs per S-haplotype |
| $E_{\text{th}}$ | Interaction energy threshold |
| $\mu = 10^{-4}$ [1/generation] | Per-residue mutation rate |
| $L = 18$ | Number of residues (amino acids) in each gene |
| $\alpha = 0.90$ | Proportion of self-pollen received by haploid maternal plant |
| $\delta = 0.90$ | Inbreeding depression – the proportion of self-fertilization inviable offspring |
| $k = N$ | The number of non-self fertilization attempts for a female S-haplotype |

**Table 3:** Compatible fraction and average number of SLFs per S-haplotype interacting with 1 RNase. The first 3 rows of the Table show the data of Fig. 4a.

| $E_{\text{th}}$ | -10 | -9 | -8 | -7 | -6 | -5 | -4 | -3 | -2 |
| --- | --- | --- | --- | --- | --- | --- | --- | --- | --- |
| Compatible fraction (neutral) | 0.032 | 0.084 | 0.182 | 0.334 | 0.515 | 0.698 | 0.84 | 0.929 | 0.975 |
| Compatible fraction (evolved) | 0.285 | 0.224 | 0.224 | 0.212 | 0.226 | 0.285 | 0.371 | 0.439 | 0.519 |
| Compatible fraction (marginal) | 0.008 | 0.027 | 0.052 | 0.107 | 0.182 | 0.278 | 0.413 | 0.54 | 0.604 |
| Average number of SLFs per S-haplotype interacting with 1 RNase (all randomly drawn from the prior distribution) | 0.25 | 0.67 | 1.49 | 2.76 | 4.37 | 6.02 | 7.37 | 8.26 | 8.72 |

Our mutation mechanism replaces amino acids at random with those drawn from the prior distribution. Hence, if mutations are repeatedly applied without any selection pressure, the marginal distributions *ψ*^RNase^(*i*), *ψ*^SLF^(*i*) should follow the prior distribution *ν*(*i*). Since mutations occur independently in the RNase and SLF, their joint AA distribution *φ* in the neutral case would be the product of these two marginals, namely *φ*(*i, k*) = *ν*(*i*) × *ν*(*k*). Thus, deviations of either *ψ* or *φ* from the prior distribution *ν* are a signature of selection pressure operating beside the mutations.

To quantify deviations of the amino acid frequencies from the prior, we employ the Kull-back–Leibler divergence *D*_KL_ (a.k.a relative entropy). *D*_KL_ is an information-theoretic measure of divergence between two distributions *P* (*x*) and *Q*(*x*), defined as [34]

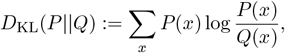

and measured in units of nats (natural bits). To identify signatures of selection we calculated *D*_KL_(*ψ*^*j*^||*ν*), *j* ∈ {RNase, SLF} the divergence of each marginal distribution from the prior *ν*(*i*) for different levels of *E*_th_ (Fig. 4b). Since a match at random between RNase and SLF is more likely under higher promiscuity, we expect the selection pressure for match avoidance to intensify with *E*_th_. Indeed, we observe a larger deviation from the prior as *E*_th_ increases for the RNase distribution *ψ*^RNase^. Yet, the SLF distribution remains close to the prior for the entire *E*_th_ range, namely *D*_KL_(*ψ*^SLF^||*ν*) ≈ 0. In the last section we provide a theoretical argument for this asymmetric response.

Compariosn between the evolved and neutral curves in Fig. 4a suggests that in most of the range (*E*_th_ *>* −8) the dominant selection pressure affecting AA frequencies is self-compatibility avoidance. To further test this hypothesis, we ran an additional simulation of a reduced model in which we only applied selection pressure to avoid self-compatibility. To do so we allowed individuals to regularly mutate, permitting reproduction without testing compatibility, and applying selection only by discarding every self-compatible mutant. We denote the marginal AA distributions obtained in the reduced model by 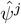, *j* ∈ {RNase, SLF}, respectively. Similar to the full model (Fig. 4b), we found that in the reduced model too, only the RNase AA frequencies 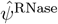were biased, but the SLF AA frequencies closely followed the prior (Fig. S12). To directly compare the two models we compute the divergence of the full simulation with respect to the reduced model (Fig. 4c) and observe that it is less than 0.05 nat per AA base, so that the two models are almost in perfect agreement.

If the selection pressure to avoid within-S-haplotype matches dominates, we also expect it to increase with the number of SLF genes per S-haplotype. Indeed, in Fig. S12 we illustrate that *D*_KL_(*ψ*^RNase^||*ν*) between the evolved and prior distributions also increases with the number of SLF genes per S-haplotype for fixed *E*_th_. Again, we observe that the RNase bears the vast majority of the selection pressure with only little impact on the SLF sequences. We also simulated selection to avoid self-compatibility alone in a symmetric setting of one RNase and only one SLF per S-haplotype (Fig. S12). Here, for the first time, we observe a symmetric response to selection, where both RNase and SLF AA distributions 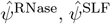are equally biased relative to the prior distribution 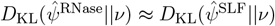.

The balance between the selection pressures to attract and repel also depends on the pollen-to-ovule ratio, represented in our model by the number of fertilization attempts *k* per female. Under pollen abundance, RNase evolution is almost entirely governed by the pressure to avoid self-compatibility. Pollen limitation (low *k*), gives rise to an additional selection pressure on the RNase to match SLF partners, while the previous pressures on SLFs to match, and on both to avoid self-compatibility, persist. Thus, pollen limitation tips the balance between the attraction and repulsion pressures. The results above were obtained under pollen abundance. We found however that only for very low values of *k* = 1, 2 the amino acid content differs, whereas for *k >* 5 it is virtually insensitive to *k* (Fig. S8).

### Matches between proteins can be implemented by either ‘global’ or ‘local’ strategies

Evolving a sequence to match or mismatch other sequences can employ either of two basic strategies (or their combination), which we term ‘global’ and ‘local’. A global strategy means that proteins are broadly attractive or repulsive regardless of the particular partner-sequence, and a local strategy means that the protein selectively matches or mismatches only particular partners. A global strategy is implemented by biasing the proein’s marginal AA frequencies in a position-independent manner, while a local strategy relies on specific position-dependent adjustments to particular partner-sequences. A local strategy can generate a significant change in the interaction energy by controlling relatively few AAs and often does not require a drastic change in their marginal frequencies. Hence, whenever (in)compatibility with respect to only one (or very few) partners is selected for, it is expected that a local strategy will dominate, while otherwise, a global strategy is preferable (see Fig. 5d for summary). In this section we distinguish between the contributions of global and local strategies to attraction and to repulsion between proteins, by analyzing separately interactions between same-S-haplotype RNase-SLF pairs and interactions between RNase-SLF partners from different S-haplotypes. To quantify the contribution of the local strategy we calculate the deviation of the joint RNase-SLF AA frequencies from the product of the marginal frequencies again by applying the Kullback-Leibler divergence.

**Figure 5:**
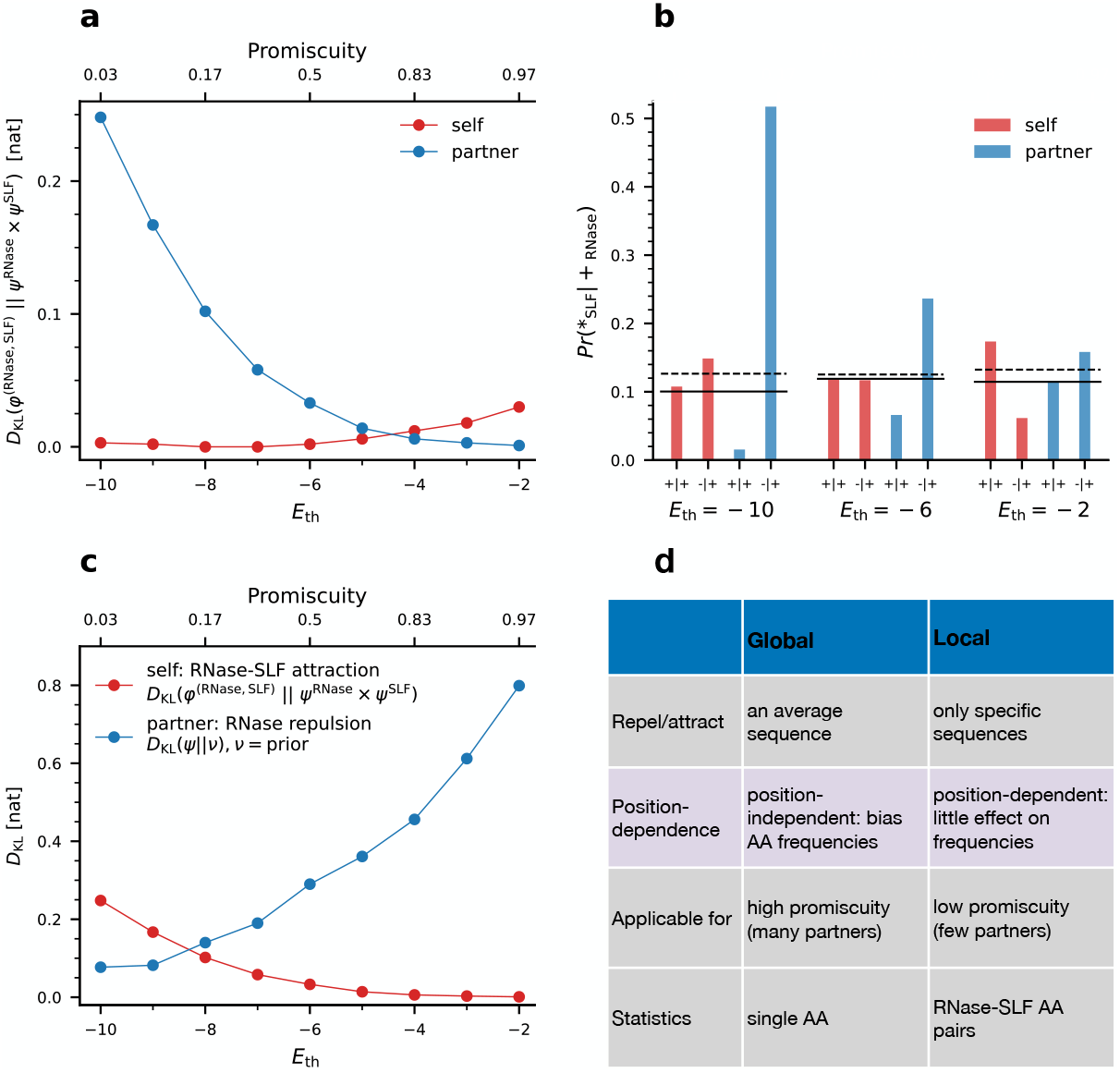
Protein pairs exhibit sequence coordination (‘local’ strategy) on top of uncoordinated biases in AA frequencies (‘global’ strategy). **(a)** We quantify the sequence coordination between RNase-SLF pairs using the Kullback-Leibler divergence between the empirical joint distribution *φ*^(RNase,SLF)^ and the product of their marginal distributions *ψ*^RNase^ × *ψ*^SLF^. We assess separately RNase-SLF pairs within the same S-haplotype (‘self’, red) and RNase-SLF partners located on distinct S-haplotypes (partner, blue) and test their dependence on the promiscuity. **(b)** To test whether a local strategy contributes to attraction or to repulsion between protein pairs we quantify the frequencies of charged amino acids that could play a role in both attraction and repulsion. We present here the frequencies of charged amino acids in SLFs conditioned on the RNase having a positively charged (+) residue in the corresponding position for RNase-SLF pairs either from the same S-haplotype (self, red) or for partners from different S-haplotypes (partner, blue) under three different *E*_th_ values. The horizontal lines show for reference the marginal SLF frequencies ((+) solid lines, and (–) dashed lines), where deviation from these marginals is a signature of a local strategy. **(c)** Comparing the intensity of the global strategy, manifested by the RNase deviation from the prior *D*_KL_(*ψ*^RNase^||*ν*) (blue curve) and the local strategy, manifested by the deviation of RNase-SLF partner pairs from the product of their marginals *D*_KL_(*φ*^(RNase,SLF)^||*ψ*^RNase^*ψ*^SLF^) (red curve) shows that in most of the *E*_th_, a global strategy has a greater contribution than a local one. **(d)** Qualitative comparison between global and local strategies.

To quantify to which extent the compatibility gap of Fig. 4a between evolved and neutral sequences is explained by biases in the single AA frequencies of the evolved sequences (‘global’), we produced alternative test sequences that were not evolved in simulation, but only had the same marginal AA distributions as the evolved ones. We calculated the compatible fraction among those fabricated test RNase-SLF pairs (Fig. 4a, gray curve, ‘marginal’). We observe that for high promiscuity, the ‘marginal’ curve is close to the ‘evolved’. We conclude that most of the compatibility reduction of the evolved sequences relative to the neutral ones is achieved by position-independent biases in AA frequencies, namely a global strategy. Yet, for low-to-intermediate promiscuity the ‘marginal’ (gray curve) deviates significantly from the evolved (red curve), suggesting that a local strategy is at play there.

As a global strategy only biases a sequence’s AA frequencies independently of its partners, it was sufficient to assess the single AA frequencies to quantify its magnitude. In contrast, a local strategy induces dependencies between same-position residues of partner and anti-partner RNase-SLF pairs, often with only a minor effect on their single AA frequencies. These local dependencies are not captured by the single sequence AA frequencies, but only by the joint distribution of sequence pairs. To quantify its contribution, we calculated the deviation of the joint RNase-SLF distribution *φ*^(RNase,SLF)^ relative to the product of its marginals, *ψ*^RNase^ × *ψ*^SLF^, using Kullback-Leibler divergence *D*_KL_(*φ*^(RNase,SLF)^||*ψ*^RNase^ × *ψ*^SLF^). If these sequences evolve only via position-independent biases in their AA frequencies, the joint RNase-SLF distribution should follow the product of their marginals, whereas any deviation of *D*_KL_ from zero is indicative of a local strategy contribution.

To disentangle the opposite effects of the two selection pressures operating simultaneously to attract (across S-haplotypes) and repel (within-S-haplotype), in Fig. 5a we calculated *D*_KL_ separately for pairs of same-S-haplotype RNases and SLFs, that are under pressure to repel each other (red curve), and pairs of partner RNase-SLF located on different S-haplotypes, that are under pressure to attract each other (blue).

Promiscuity affects these two selection pressures in two ways: firstly, as *E*_th_ increases, the pressure to repel (within the S-haplotype) rises, while the pressure to attract (across S-haplotypes) lessens. Secondly, the number of partners per protein increases with promiscuity (Fig. 3b) rendering a local strategy less useful, as it is difficult to match multiple distinct partners via local sequence arrangements. Only for the lowest promiscuity, most proteins have only one partner (or none), making a local strategy feasible. In contrast, the number of anti-partners each protein should repel is the number of SLF genes per S-haplotype – a fixed parameter in our model independent of *E*_th_. Indeed, we found that a local strategy was dominant for partner attraction (Fig. 5a, blue) under low promiscuity and its contribution decreased with *E*_th_. For within-S-haplotype repulsion, we found an opposite trend, where the contribution of a local strategy to repulsion increased with promiscuity (Fig. 5a, red). Comparing the magnitudes of local strategy contributions to attraction and to repulsion, the former obviously dominates in most of the *E*_th_ range.

As the Kullback-Leibler divergence is only indicative of a deviation from the marginal distribution but not of its directionality, we further refined our analysis to distinguish between the opposite effects of repulsion and attraction. In Fig. 5b we present the conditional RNase-SLF AA frequencies of charged AA distinguishing between same-charge (repulsive) and opposite charge (attractive) AA pairs, compared to the marginal frequencies (horizontal lines). Again, we look separately at pairs of same-S-haplotype RNase and SLF evolving to repel each other (red) and at pairs of RNase and SLF partners, evolving to attract each other (blue).

In line with the dominance of a local strategy under low *E*_th_ as demonstrated in Fig. 5a, we find among partner RNase-SLF pairs, an over-representation of +/- (attractive) and under-representation of +/+ (repulsive) AA pairs, suggesting a dominant role for a local strategy in partner attraction. The magnitude of this effect is highest under low *E*_th_ (few partners) and decreases with *E*_th_ as the typical number of partners per protein increases and a local strategy is useless (Fig. 5b). We find qualitatively similar trends for H-H and P-P AA pairs that also promote attraction (Fig. S7).

In contrast, for same-S-haplotype RNase-SLF pairs that evolve to repel each other (Fig. 5b, red bars) we find much smaller deviations from the marginals, showing only little contribution of a local strategy to avoidance of self-compatibility. This result too is in agreement with Fig. 5a demonstrating a lesser role for a local strategy among same-S-haplotype pairs.

### The amino acid frequencies evolve to minimize the divergence from the prior distribution under the constraint of no self-compatibility

We found that the dominant selection pressure governing AA composition is avoidance of self-compatibility, but why would it act almost exclusively on the RNases with only little effect on the SLFs (Fig. 4b)? At first glance, this seems non-intuitive, given that under pollen abundance as simulated here, selection to match acts mostly on the SLFs and much less on the RNases receiving multiple fertilization attempts each. Thus, one may naively expect that the SLFs should be the ones changing their AA content in response to selection, whereas the RNases will evolve almost neutrally. Below we introduce a theoretical argument using large deviations theory to establish that self-compatibility avoidance alone well-approximates the amino acid frequencies in the S-haplotype and explains the asymmetric response to selection of the RNase and SLF.

Recall, that in the absence of selection or any other constraint, when the process is solely governed by mutations drawn from the prior distribution *ν*(*i*), the stationary pairwise AA distribution *φ*(*i, k*) follows *φ*(*i, k*) = *ν*(*i*) × *ν*(*k*). We analyze this system subject only to the constraint of self-incompatibility elimination, as in the reduced model introduced above. In this section, we provide only a sketch of this analysis, while more rigorous arguments are postponed to a matematical appendix.

Fixing *E*_th_, we analyze the stationary joint RNase-SLF AA distribution via energy-entropy considerations, applying large deviations theory. A balance between two forces determines the simulated AA distribution. The first is the entropy of the system, namely the number of sequences with a given AA distribution. The second is the long term survival probability of a sequence, here equivalent to the long term probability that mutations will not induce self-compatibility. Large deviations theory tells us that for such systems, when *L*, the sequence length, tends to infinity, the AA composition will be close, with high probability, to the distribution which maximizes the product of the number of configurations and the survival probability. This phenomenon is known as the energy-entropy maximization principle [35]. While this product is analytically intractable, we can well approximate the survival probability by a monotone function of the energy of the system, as RNase-SLF pairs having a higher interaction energy require a larger number of mutations, before they become self-compatible. In such a setting, the distribution maximizing the energy-entropy product could also be well approximated as the distribution which maximizes only the entropy, subject to a harsher effective energy threshold which we denote by 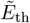. More concretely, it is well approximated by the survival distribution in a simplified model with 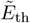, where haplotyes are eliminated only if they are self-compatible at the end of the simulation. A decoupling argument justifying this approximation appears towards the end of the appendix.

For such a simplified model, Sanov’s Theorem [35], a classical tool from large deviations theory, enables us to analytically compute an excellent approximation of the final single and pair marginals of the AA distribution, as the one which minimizes the divergence with respect to AA independently drawn according to *ν*, among all those which satisfy the non-self-compatibility condition (see Eq. (9) in the appendix). They also tell us that these marginals determine the entire AA distribution of the S-haplotype. This derivation is detailed in the first part of the appendix.

These predictions should be valid regardless of the inbreeding depression *δ*, which is only accounted for by altering the effective energy threshold 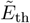(see appendix). This approximation should persist as long as avoiding self-compatibility is the main evolutionary pressure at work, indicating that in this regime AA frequencies should depend smoothly on *E*_th_ and *δ* (see Fig. S6). In Fig. 6 we compare the AA frequencies obtained in the simulations of the full and reduced models to the prediction obtained by numerical minimization of Eq. (9) for the entire range of *E*_th_ simulated. Except for the lowest promiscuity, the two models yield very similar AA frequencies, and Eq. (9) provides an excellent approximation to the reduced model. Both the full and the reduced models predict a depletion of hydrophobic and an enrichment of charged AAs in the RNase, where these trends increase with *E*_th_. We compared these theoretical predictions to the AA frequencies of RNase residues computationally estimated as being under positive selection[19]and likely serving as specificity-determinants (Fig. 6, shaded region in each panel). In this empirical data too we find a similar trend with under-representation of hydrophobic and over-representation of charged AAs, in agreement with our theoretical prediction.

**Figure 6:**
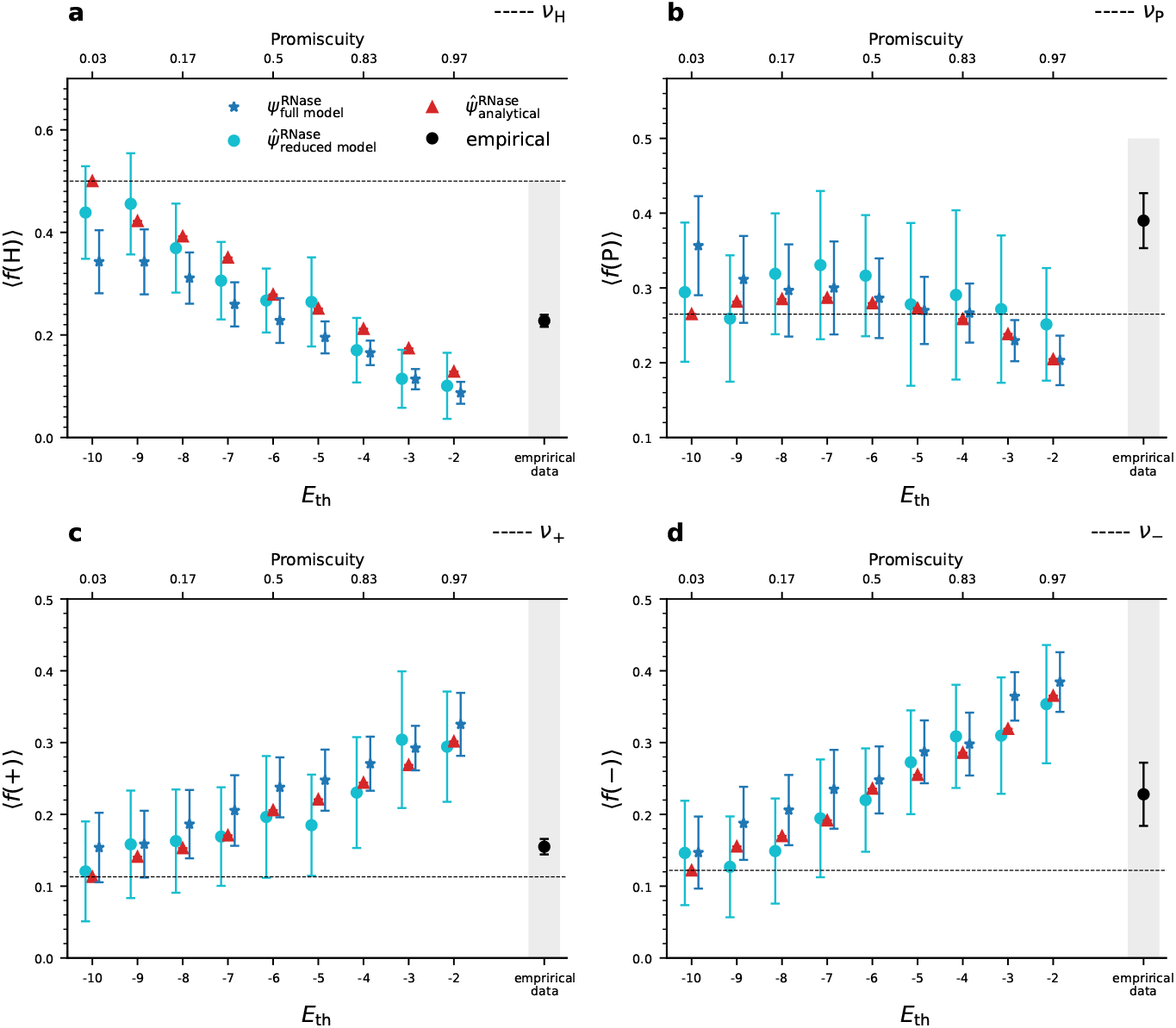
The marginal AA frequencies in RNases in the full model and reduced model simulations, are in good agreement with the analytical approximation; Empirical data shows a similar trend to our theoretical predictions. We plot the RNase frequencies of the four AA categories (H, P, +, –) in the full model (blue star) and reduced model (cyan circle) simulations, and the analytical approximation (red triangle) for different *E*_th_ values with 9 SLFs per S-haplotype. The horizontal dashed lines mark the prior frequencies of the corresponding AA categories obtained under neutrality. We find an decrease in the frequency of hydrophobic and increase in the frequncies of charged AA with *E*_th_, in agreement with the increasing pressure to avoid self-compatibility. The black circle (shaded background) in each panel shows the frequency of the corresponding AA category in empirical data of RNase residues computationally estimated to be under positive selection [19] (Methods) exhibiting a similar trend with under representation of hydrophobic and over-representation of charged AA relative to neutrality. Symbols denote averages over multiple simulation data points, and bars represent ±STD relative to the mean. The figure is based on 25 independent runs for each *E*_th_ value, 450 data points taken from each run, with a 25-generation interval between consecutive time points. All these data points were taken after the entire population descended from a single common ancestor and the amino acid frequencies had already reached their steady-state proportions. The total runtime for each *E*_th_ value is 1.5 × 10^5^ generations. Parameter values are as in Table 2.

To understand the asymmetric response to selection of the RNases and the SLFs, observe that in Eq. (9) every modification in the frequencies of the SLFs is *m* times as costly as its counterpart for the RNase. This is because a single RNase has to avoid compatibility with *m* SLFs. As a result, to minimize the entropic cost of avoiding self-compatibility, the RNase AA distribution should diverge from the prior distribution significantly more than the SLF AA distribution should. The full simulation sees an additional pressure on the SLFs to match their RNase partners.

While this has little effect on the behavior of the RNases, it could, in principle, tilt the AA frequencies of the SLFs. The fact that this additional pressure does not alter the joint distribution *φ*^(RNase,SLF)^ in the full model simulations is explained by the use of a local rather than a global strategy to match.

## Conclusions

Combining the results of Figs. 4-6, we conclude that under most of the parameter regime, there are many-to-many matches where proteins evolve by tuning their attractiveness/repulsion towards a generic partner and do so by biasing their amino acid frequencies to deviate from the prior ones (global strategy). This strategy allows to easily match newly formed proteins, facilitating the emergence of new classes. Only in the extreme of low promiscuity, the proteins mostly have only a single partner (or none), hence the strategy of tailoring their sequence to match a specific partner becomes useful. This local strategy is mostly used for partner attraction and has only a minor role in anti-partner repulsion. See Fig. 5c for comparison between the magnitudes of global and local strategies in the entire *E*_th_ range.

Roughly speaking, RNases develop non-specific locks with calibrated difficulty to guarantee self-incompatibility (global strategy, Fig. 4a). The SLFs, in turn, develop tailored keys to unlock particular RNases (local strategy), so that each paternal S-haplotype can fertilize maternal S-haplotypes with nearly all non-self RNases. As *E*_th_ decreases, the SLFs specialize in unlocking a decreasing set of RNase locks.

This also explains why the reduced model, in which the selection to match was not accounted for, provides a good approximation to the frequencies in the full model: when promiscuity is high, the selection pressure on the SLFs to match is minor, as they have a high probability of matching at random, and when promiscuity is lower, the selection pressure is expressed by a local matching strategy, which has only a minor effect on the AA frequencies in a single S-haplotype.

## Discussion

The collaborative non-self recognition self-incompatibility system in hermaphroditic plants is an example of a complex biological system comprised of multiple components that co-evolve under a set of potentially conflicting selection pressures. The evolutionary unit in this system is an S-haplotype that includes one female (RNase) and multiple male (SLFs) type-specifying genes that are tightly linked and consequently affect each other’s evolution. We recently proposed a theoretical framework that incorporates molecular recognition between the male and female type-specifying proteins into the evolutionary model [33]. Our model builds on the cornerstones of a degenerate genotype-to-phenotype mapping allowing multiple genotypes to share a common compatibility phenotype and on the promiscuity of molecular interactions. Using this model we study how its numerous microscopic degrees of freedom (in the single gene and in the individual levels) robustly give rise to a few population-level macroscopic phenotypes. Our model does not include an explicit definition of fitness as often employed in evolutionary models. Instead, an individual’s reproductive success is determined by the interaction energies amongst its self-proteins and between its proteins and those of other population members, inducing dependencies of an individual fitness on the entire population composition. Matches and mismatches between individuals establishing the macroscopic system states can be achieved by a variety of amino acid (AA) sequences constituting its microscopic states. Hence, our model exhibits a degenerate genotype-to-phenotype mapping.

In plant populations, pollen is often more abundant than ovules. Hence, many pollen grains compete over fewer ovules resulting in asymmetry between the selection pressures acting on the hermaphroditic plant in its roles as male and as female. Thus, the pressure for cross-compatibility in the collaborative non-self recognition system is mostly borne by the SLF (male) and much less by the RNase (female) genes. In contrast, the pressure to avoid self-compatibility is shared by both, but is much more expressed in the RNase which has to avoid match with multiple SLFs. If selection were only to match, one could have expected the RNases to employ the neutral AA frequencies, while the SLF frequencies were biased by the pressure to match. However, because of the additional and dominant pressure to avoid within-S-haplotype match, we observe a very different response, where the SLF frequencies are neutral and the RNases are pushed to avoid match and hence their frequencies deviate from neutrality. While the magnitude of both selection pressures varies with promiscuity, the pollen-to-ovule ratio and the inbreeding depression, the dominance of selection to avoid self-compatibility holds under a broad range of these parameter values (Fig. 4, Fig. S8), and it is only reversed in the non-promiscuous region *E*_th_ ≤ −9 or under severe pollen limitation. In the latter case, the weight of the selection pressure to match acting on the RNases increases as compatible pollen becomes scarce, while the pressure to avoid self-compatibility remains intact, thus tipping the balance between these two pressures acting on the RNases (Fig. S8).

The asymmetry in response to selection between the RNases and SLFs is explained by an entropic argument: due to the multiplicity of SLFs, biasing their AA frequencies bears a larger cost of entropy reduction compared to the cost required to bias the RNase, that has only a single copy per S-haplotype. We frame this argument using large deviations theory, whose accuracy improves as the sequence length *L* increases. While we use here *L* = 18, in accordance with the typical size of protein binding domains and typical numbers of residues estimated to be under positive selection, we obtain a very good fit between the theoretical prediction and the simulation results. The balance between multitude and fitness (reminiscent of the energy-entropy balance in statistical physics) can be used to infer the statistics of biological systems exhibiting a degenerate mapping between genotype to phenotype. Another example for a system in which similar considerations were applied to infer the stationary distribution is transcription factor binding sites [36, 37, 30]. Similar to the self-incompatibility system, there too the phenotype is determined by the binding energy alone and can be achieved by a multitude of distinct sequences. We found that in the promiscuous region, where the system is qualitatively compatible with empirical data, evolved sequences match each other much less than neutral ones do. The default being high compatibility between proteins, unless selection operates to reduce it, and the dominance of selection against avoid self-compatibility are counter-intuitive. These observations are in line with an experimental study by Zhao *et al*. [38] that demonstrated 85% compatibility between two Petunia RNases and several SLFs of different species or families, which obviously cannot fertilize Petunia. In contrast, only 19% of same-species SLFs exhibited such compatibility. In light of our model, this empirical result can be interpreted as compatibility being the default between RNase and SLF proteins in the absence of selection, as is the case for different species that cannot fertilize each other. Compatibility decreases only amongst same-species proteins for which selection against self-compatibility operates. This empirical result suggests though that local matches play a more dominant role in AA frequencies in the elimination of self-compatibility than global biases do, whereas our model predicts that elimination is almost exclusively achieved by a global change in AA frequencies.

A decrease in hydrophobicity as a means for a protein to be indiscriminately less attractive, as predicted by our model and corroborated by empirical RNase data (Fig. 6), was previously reported as a transient state when proteins switch between different partners. Likewise, an increase in charged amino acid content was proposed to later follow as a means to specifically tune to attract or repel certain proteins [39]. Assessment of the amino acid content of hub proteins also showed a decrease in hydrophobic and an increase in electrostatic interactions compared to no-hub proteins [40], suggesting that these features are more broadly applicable.

A simplifying assumption in our model is that the proteins bind each other in one orientation only. It remains unknown whether SLFs having multiple RNase partners use a single shared binding domain for all partners or alternatively have a separate interface dedicated to each [40]. It would be interesting to extend our model to allow for multiple binding geometries and study how this affects the balance between global and local match strategies and the resultant amino acid frequencies.

Our model exhibits several key microscopic properties that give rise to its macroscopic behavior: promiscuity, degenerate mapping between genotype-to-phenotype, and a co-evolution of multiple components under a combination of positive and negative design pressures. These properties are not unique to the self-incompatibility system and were observed in several other biological systems. For example, a degenerate genotype to phenotype mapping is known in transcription factor-DNA binding [30] and RNA secondary structures [27]. Co-evolution of components that should match certain partners (‘positive design’) but avoid others (‘negative desiogn’) is known in a variety of biological systems, such as two-component signaling systems [41], bacterial toxin-antitoxin [42], host-pathogen [43], and venoms [44, 45], and are often accompanied by selection for diversification [46, 43, 42]. The superposition of positive and negative designs [47, 48] also plays a key role in the evolution of the immune system whose aim is to distinguish between self and non-self peptides [49] and in gene regulatory networks where transcription factors should distinguish between cognate and non-cognate binding sites [50]. Promiscuity in molecular recognition was observed in signaling proteins [51], and in transcription factor-DNA binding [52] and proposed to be pivotal in the evolvability of gene regulatory networks [53, 54] and in the immune receptor functionality [55, 56]. Following these commonalities, we propose that the incorporation of molecular recognition into the evolutionary model may be more broadly applicable and further ramifications of our framework could be exploited to study additional systems. It would be interesting to see whether these systems also share some of the behaviors exhibited by the plant collaborative self-incompatibility system, as shown here.

## Methods

### Four biochemical categories of amino acids (AA)

We have classified the 20 AAs into four different biochemical categories: hydrophobic (H), neutrally polarized (P), positively charged (+), and negatively charged (–), see Table S1. The prior frequencies of the different categories are given by *ν*(*i*|*i* = {H, P, +, −}) = {0.5, 0.265, 0.113, 0.122}, respectively [33]. Table 1 shows the interaction energies between pairs of amino acids of all possible categories. These values were adopted from [57].

### Parameters and default values

The model parameters and their default values (unless specified otherwise) are summarized in Table 2. The parameter *E*_th_, is the main parameters that vary from -10 to -2 with step size 1.

### The stochastic simulation

#### Initialization of the S-haplotype population

We initialize the S-haplotype population using *n* distinct types, such that each type is self-incompatible and bidirectionally compatible with the remaining *n* − 1 types. We initialize the population with equal frequencies of all types. We employ the following three steps to draw these initial types:

1. **Draw RNase sequences**: First, generate *n* distinct RNases R = {R_1_ … R_*n*_} by randomly drawing sequences from the prior distribution *ν*.
2. **Draw SLF sequences**: Construct *n* pools where each pool P_*i*_ = {S_*m*_}, *i* ∈ (1 … *n*), contains *m* SLF sequences, that can detoxify the *i*-th RNase, and potentially others as well. To construct the SLF pools, we generate SLFs, one at a time, test which RNase(s) it detoxifies, and add that SLF into the corresponding pool(s). Any SLF that does not detoxify any RNase or that detoxifies all of them is discarded. Repeat this step until each pool has *m* SLF genes.
3. **Form** *N* **self-incompatible and complete S-haplotypes**: For each RNase R_*i*_ ∈ R, choose in total *n* − 1 SLF genes, one from each pool P_*j*_, *j* ∈ (1 … *n*), *j* ≠ *i* and make sure none of these SLFs detoxifies R_*i*_. If it does, replace it with another SLF from the same pool until an SLF that does not detoxify R_*i*_ is found. After generating *n* distinct S-haplotypes, we construct an initial population of size *N* using exactly *N/n* copies of each S-haplotype, and then shuffle them.

The above procedure produces SLFs that can potentially match multiple RNases each. Alternatively, to construct an initial population with only one-to-one RNase-SLF interactions, each pool P_*i*_ should have only those SLFs that detoxify only R_*i*_.

#### Life-cycle of the full model simulation

1. **Mutation**: Each amino acid in each gene can mutate with probability *µ* per generation. An amino acid chosen to mutate is replaced by a randomly chosen amino acid drawn from the prior distribution *ν*.
2. **Maternal S-haplotype picking**: Maternal SC S-haplotypes can be fertilized by either non-self paternal S-haplotype or by the self one. In the latter case, only a proportion 1 − *δ* of the offspring produced via self-fertilization survives. Maternal S-haplotypes chosen to reproduce can be either self-compatible (SC) with either self or non-self fertilization or self-incompatible (SI) allowing only non-self fertilization. Each generation we calculate the probabilities of either group forming offspring:

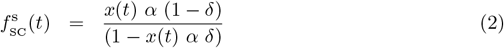

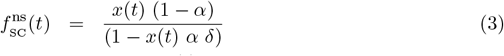

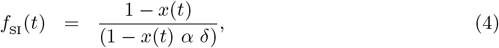

where *x*(*t*), and 1−*x*(*t*) are the population fraction of SC and SI S-haplotypes in the current generation, *α* is the proportion of self-pollen received and *δ* is the inbreeding depression, i.e. the proportion of self-fertilization offspring that do not survive. Full derivation of 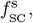 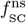, and *f*_SI_ is provided in the supplementary information file.

**3.Offspring formation**: Pick a maternal S-haplotype H_*i*_ from either group with probabilities 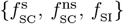. If the maternal S-haplotype H_*i*_ belongs to 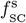, the offspring is identical to the parental one. If the maternal S-haplotype H_*i*_ belongs to either 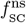 or *f*_SI_, it is fertilized by non-self pollen. In this case, it is given *k* opportunities to match a randomly chosen non-self S-haplotype partner, in the paternal role. If such a match is found, the offspring would be one of the two parental S-haplotypes with equal probability. Repeat this step until *N* offspring are formed.

Steps 1-3 constitute a single generation. Repeat these steps until the desired number of generations is reached.

Total simulation run times for the default parameter values are listed in Table 4. The analyses are usually based on a total of 25 independent runs for each parameter set, where 1200 data points are taken from each run, with a 25-generation interval between consecutive time points.

**Table 4:** Typical simulation times for different *E*_th_ values.

| $E_{\text{th}}$ | -10 | -9 | -8 | -7 | -6 | -5 | -4 | -3 | -2 |
| --- | --- | --- | --- | --- | --- | --- | --- | --- | --- |
| Generations ($\times 10^5$) | 1.5 | 1.5 | 1.5 | 1.5 | 2.0 | 2.75 | 4.5 | 4.5 | 4.5 |

### Life-cycle of the reduced model simulation

1. **Mutation**: Each amino acid in each gene can mutate with probability *µ* per generation. An amino acid chosen to mutate is replaced by a randomly chosen amino acid drawn from the prior distribution *ν*.
2. **Removal of self-compatible S-haplotypes** The previous mutation can potentially create self-compatible S-haplotypes. We discard these self-compatible S-haplotypes, if they exist, and remain with only *Ñ* ≤ *N* self-incompatible ones.
3. **Parental S-haplotype picking**: We randomly pick *N* parental S-haplotypes with replacements from the remaining *Ñ* self-incompatible ones. The chosen S-haplotypes will form the offspring population.

Here steps 1-3 form a single generation.

### Simulation run-time

We simulated the model for different numbers of generations as described in Table 4. The two criteria should be met to stop the simulation: the amino acid frequencies of RNases and SLFs should stabilize and the entire population should be the descendants of only one of the *n* initial S-haplotypes. For *E*_th_ = -2, it takes a larger number of generations to achieve the latter, compared to the former criterion.

### Simulation data analysis

#### RNase partners per SLF gene

For a given run *j*, and at generation *t*, consider 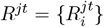 to be the set of all unique RNases, where 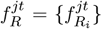 is the set of their corresponding frequencies. Similarly, let 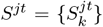, and 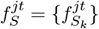be the set of all unique SLFs and their corresponding frequencies. Further, consider 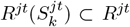 to be the set containing all distinct RNases that are detoxified by a particular SLF gene 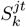. The effective number of RNase partners of SLF gene 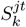is given by:

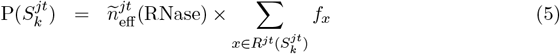

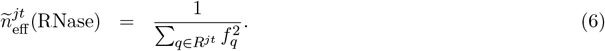

where *f*_*x*_ is the population frequency of unique RNase *x*.

### Distribution of the number of partners per SLF

Let 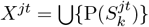 be the set of unique partner numbers of all unique SLF genes. For a fixed bin size *h*, and maximum number of RNases *M* over all independent runs and for all data points a fraction of SLF genes *x*^*S*^ detoxifying a number of RNases in the range [*i h*, (*i* + 1) *h*), *i* ∈ (0, *M*) is calculated as,

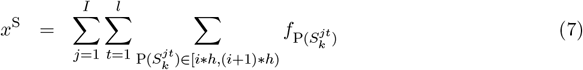

where 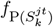is the frequency of the *k*-th SLF gene that has a number of partners 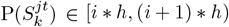. In Fig. 3a,b we took *h* = 1.0.

where *f*_*x*_ is the population frequency of unique RNase *x*.

### The effective energy threshold used in the analytical approximation

In the analytical calculation, we considered an effective energy threshold 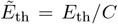, where the value of *C* was extracted from our simulations of the reduced model (Fig. S9), as the ratio of the *E*_th_ value used in simulation and the average of total interaction energy obtained in evolved populations, 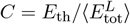, thus requiring a harsher energy threshold to avoid self-compatibility than used in the simulations (see appendix for more details). See supplementary Mathematica file for the numerical calculation.

### Amino acid composition in RNase empirical data

The empirical data points in Fig. S2 are based on the positively selected positions in RNases of *Petunia inflata, Petunia hybrida*, and *Petunia axillaris* as inferred computationally by [19]. We classify the amino acids in these positions to either of the four biochemical classes based on the classification in Table S1.

### Mathematical Appendix

To establish the fact that avoiding self-compatibility is the primary evolutionary force governing the evolution of the amino acid frequencies in the S-haplotype and explain the asymmetric response to the selection of the RNase and SLF, we estimate the simulated amino-acid distribution in an S-haplotype, evolved strictly under pressure to avoid self-compatibility. We then show that this distribution is nearly identical to the distribution in our full model simulation, by comparing the relative AA frequencies. To complement this empirical computation, we also consider a theoretical estimate derived in two steps.

#### Step 1: Analyzing a simpler model governed only by entropy considerations

Firstly we consider an even simpler non-evolutionary model. Given an S-haplotype, denote by *E*_*i*_ its interaction energy between the RNase and the *i*-th SLF as per Eq. (1). In the simplified model we pick at random a neutral S-haplotype with a single RNase and *m* SLFs, each of length *L*, drawn from the prior distribution, conditioned on *E*_*i*_ *> E*_th_ for all *i*, where *E*_*i*_ is the interaction energy between the RNase and the *i*-th SLF. Observe that this corresponds to a situation in which the population evolves neutrally and then suddenly every self-compatible S-haplotype is eliminated, and we are interested in the typical distribution of a surviving S-haplotype.

To understand the behavior of such a sample, assuming that *L* is sufficiently large, we may apply Sanov’s Theorem, which tells us that, as the sequence length *L* tends to infinity, the sampled distribution conditioned to be in any non-empty set Γ will be close to the minimizer of the relative entropy with respect to the prior distribution.

Let us state this formally. Let *q* be a probability distribution on an alphabet Σ, and write 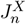for the AA-frequencies of a sequence *X* = (*X*_1_, …, *X*_*n*_) ∈ Σ^*n*^ whose coordinates are drawn independently according to *q* (in our case the simulated S-haplotypes amino acids’ distribution). Consider a restricted AA-frequencies space Γ (in our case those for which the interaction energy between the RNase and every SLF is lower than the threshold). Then

**Theorem 1 (Sanov ‘57 [35][Theorem 2.1.10])**

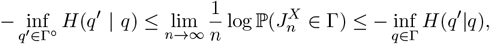

where *H*(*q*^*′*^|*q*) is the relative entropy of *q* with respect to *q*^*′*^ given by

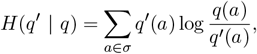

and Γ^*°*^ is the interior of Γ with respect to the weak topology. Moreover, the empirical distribution is, with high probability, in any neighborhood of the minimizers of this relative entropy.

Applying this to our model, our alphabeth consists of *n* = *L*(*m* + 1) residues, each being one of {+, −, *H, P*}. We take Γ to be the set of such distributions whose total energy between every SLF and the RNase 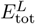is greater than *E*_th_. For large sequence length *L* we obtain that

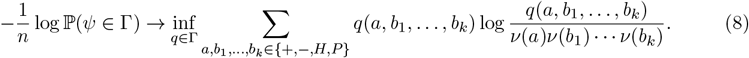

By convexity and symmetry arguments, we may restrict ourselves to *q* obtained through pair marginals between the RNase AA and each SLF AA at the same sequence position. Namely, such that there exist *ψ*^RNase^, and *φ* where *q*’s RNase AA is drawn according to the marginal *ψ*^RNase^(*a*) = _*b∈*+,*−,P,H*_*φ*(*a, b*), and, conditioned on the RNase AA *a*, each of its SLF AAs *b*_1_, …, *b*_*m*_ are drawn independently according to the conditional distribution 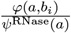.

Restricting such *q* to Γ is equivalent to restricting *q* to have marginals (*ψ*^RNase^, *φ*) for each of the *m* pairs (*a, b*_*i*_), within the family

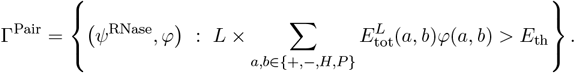

Hence we reduce the right-hand-side of Eq. (8) to

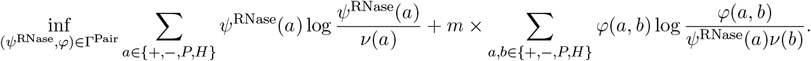

We now apply additional convexity and symmetry arguments (namely that the optimizer must be obtained when, conditioned on the RNase locus, all the SLF distributions for that locus must be identical). Hence it would suffice to seek an optimizer *q* for which there exist *ψ*^RNase^, and *φ*^(RNase,SLF)^ such that *q*’s RNase AA are drawn according to the marginal *ψ*^RNase^(*a*) = Σ _*b∈*+,*−,P,H*_ *φ*^(RNase,SLF)^(*a, b*), and, conditioned on the RNase AA *a*, each of its SLF AAs *b*_1_, …, *b*_*m*_ are drawn independently according to the conditional distribution 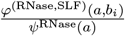. Formally, we seek such and express this in terms of *ψ* and *φ*, as minimizing

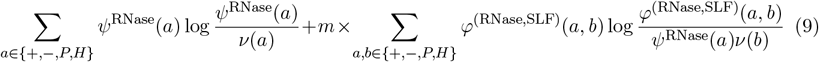

under the constraint that *ψ*^RNase^ is the marginal of *φ*. Such a minimizer could be easily recovered analytically.

#### Step 2: Relating the entropic model, to the reduced evolutionary model

Next, let us explain how to use this analysis for a model closer to our simulated reduced evolutionary model, governed only by avoiding self-compatibility. In the model analysed each S-haplotype undergoes neutral evolution, but if at any point in the course of its evolution it crosses the energy threshold – it is eliminated.

Formally, this allows us to rewrite the survival probability as P(*ψ* ∈ Γ) as

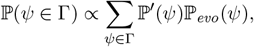

where ℙ^*′*^(*ψ*) is the neutral probability of obtaining *ψ* as in the simplified non-evolutionary model, and ℙ _*evo*_(*ψ*) is the probability of surviving above the threshold throughout the evolution, conditioned on achieving final AA-frequencies *ψ*. In the special case that ℙ _*evo*_(*ψ*) = *f* (*E*_tot_(*ψ*)) for some monotone decreasing function *f*, we can express this sum as:

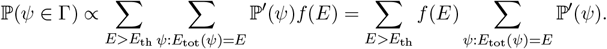

Writing 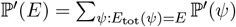 we obtain

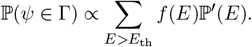

This sum has polynomially many terms, so that we may obtain a polynomially tight estimate of ℙ (*ψ* ∈ Γ) as its maximizer, namely:

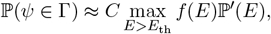

for some constant *C*. Denoting by *E*_max_ the energy maximizing this expression, we may further write

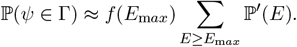

We may now apply Sanov’s theorem to the set Γ = {*ψ* : *E*_tot_(*ψ*) ≥ *E*_m*ax*_} to recover the asymptotic decay of this probability and the limiting empirical distribution, which is equivalent to an application of the standard Sanov’s theorem with a harsher energy threshold 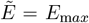. This justifies our usage of Sanov’s theorem with an effective energy threshold as per Figure S9. Observe that *δ*, the probability that an offspring formed by self-fertilization does not survive, affects this analysis only by altering the effective threshold 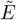, as long as the assumption that survival is roughly a function of the energy holds.

It turns out that analyzing this simpler model in which the population members are statistically independent, and the assumption that survivability is energy dependent, is sufficient to provide a good approximation of the frequencies obtained in the simulated evolutionary model. Indeed, we have formally derived the balance between the RNase and SLF amino acid frequencies and solved it numerically for the case of only selection to avoid self-compatibility. We then optimized it on the effective energy threshold and found excellent agreement with our simulation results (suppl. Mathematica file).

## Acknowledgements

This research was supported by the Israel Science Foundation, grant number 1889/19 (T.F.).

## Author contributions

T.F. conceptualized the project, acquired funding and supervised. A.J. wrote and ran the simulation, collected and analyzed results and prepared figures. K.E wrote an earlier version of the simulation. O.F. contributed to the mathematical analysis. T.F and O.F developed the methodology and wrote the paper.

## Data availability

The authors declare that all data needed to evaluate the conclusions in the paper are present in the paper and/or the Supplementary Materials.

## Code availability

The code will be made available on a public repository upon acceptance for publication.

## References

1. East, E. M. & Mangelsdorf, A. J. A New Interpretation of the Hereditary Behavior of Self-Sterile Plants. Proceedings of the National Academy of Sciences 11, 166–171 (1925). URL https://www.pnas.org/doi/abs/10.1073/pnas.11.2.166. Publisher: Proceedings of the National Academy of Sciences.

2. Wright, S. The distribution of self-sterility alleles in populations. Genetics 24, 538–552 (1939).

3. de Nettancourt, D. The Genetic Basis of Self-Incompatibility. In de Nettancourt, D. (ed.) Incompatibility in Angiosperms, Monographs on Theoretical and Applied Genetics, 28–57 (Springer, Berlin, Heidelberg, 1977). URL 10.1007/978-3-662-12051-4_2.

4. Igic, B., Lande, R. & Kohn, J. Loss of Self-Incompatibility and Its Evolutionary Consequences. International Journal of Plant Sciences 169, 93–104 (2008). URL http://www.journals.uchicago.edu/doi/abs/10.1086/523362.

5. Takayama, S. & Isogai, A. Self-Incompatibility in Plants. Annual Review of Plant Biology 56, 467–489 (2005). URL http://dx.doi.org/10.1146/annurev.arplant.56.032604.144249.

6. Fujii, S., Kubo, K.-i. & Takayama, S. Non-self- and self-recognition models in plant self-incompatibility. Nature Plants 2, 16130 (2016). URL http://www.nature.com/articles/nplants2016130.

7. Shimizu, K. K. & Tsuchimatsu, T. Evolution of Selfing: Recurrent Patterns in Molecular Adaptation. Annual Review of Ecology, Evolution, and Systematics 46, 593–622 (2015). URL 10.1146/annurev-ecolsys-112414-054249.

8. Iwano, M. & Takayama, S. Self/non-self discrimination in angiosperm self-incompatibility. Current Opinion in Plant Biology 15, 78–83 (2012). URL http://www.sciencedirect.com/science/article/pii/S1369526611001452.

9. Gervais, C., Awad, D. A., Roze, D., Castric, V. & Billiard, S. Genetic Architecture of Inbreeding Depression and the Maintenance of Gametophytic Self-Incompatibility. Evolution 68, 3317–3324 (2014). URL https://onlinelibrary.wiley.com/doi/abs/10.1111/evo.12495.

10. Williams, J. S., Wu, L., Li, S., Sun, P. & Kao, T.-H. Insight into S-RNase-based self-incompatibility in Petunia: recent findings and future directions. Frontiers in Plant Science 6 (2015). URL https://www.frontiersin.org/articles/10.3389/fpls.2015.00041.

11. McClure, B. A. et al. Style self-incompatibility gene products of Nicotlana alata are ribonucleases. Nature 342, 955–957 (1989). URL https://www.nature.com/articles/342955a0. xNumber: 6252 Publisher: Nature Publishing Group.

12. Lai, Z. et al. An F-box gene linked to the self-incompatibility (S) locus of Antirrhinum is expressed specifically in pollen and tapetum. Plant Molecular Biology 50, 29–41 (2002). URL https://link.springer.com/article/10.1023/A:1016050018779.

13. Sijacic, P., Wang, X., Skirpan, A. L., Wang, Y. & al, e. Identification of the pollen determinant of S-RNase-mediated self-incompatibility. Nature; London 429, 302–5 (2004). URL https://search.proquest.com/docview/204530198/abstract/9ABEA5AF71D048EAPQ/1.

14. Kubo, K.-i. et al. Collaborative Non-Self Recognition System in S-RNase–Based Self-Incompatibility. Science 330, 796–799 (2010). URL http://science.sciencemag.org/content/330/6005/796.

15. Kubo, K.-i. et al. Gene duplication and genetic exchange drive the evolution of S-RNase-based self-incompatibility in Petunia. Nature Plants 1, 14005 (2015). URL https://www.nature.com/articles/nplants20145/fig_tab.

16. Vieira, J., Morales-Hojas, R., Santos, R. A. M. & Vieira, C. P. Different Positively Selected Sites at the Gametophytic Self-Incompatibility Pistil S-RNase Gene in the Solanaceae and Rosaceae (Prunus, Pyrus, and Malus). Journal of Molecular Evolution 65, 175–185 (2007). URL https://link.springer.com/article/10.1007/s00239-006-0285-6.

17. Vieira, J., Ferreira, P. G., Aguiar, B., Fonseca, N. A. & Vieira, C. P. Evolutionary patterns at the RNase based gametophytic self - incompatibility system in two divergent Rosaceae groups (Maloideae and Prunus). BMC Evolutionary Biology 10, 200 (2010). URL 10.1186/1471-2148-10-200.

18. Pratas, M. I. et al. Inferences on specificity recognition at the Malus×domestica gametophytic self-incompatibility system. Scientific Reports 8, 1717 (2018). URL.0.

19. Vieira, J. et al. Predicting Specificities Under the Non-self Gametophytic Self-Incompatibility Recognition Model. Frontiers in Plant Science 10 (2019). URL https://www.frontiersin.org/articles/10.3389/fpls.2019.00879/full.

20. Aguiar, B. et al. Patterns of evolution at the gametophytic self-incompatibility Sorbus aucuparia (Pyrinae) S pollen genes support the non-self recognition by multiple factors model. Journal of Experimental Botany 64, 2423–2434 (2013). URL https://academic.oup.com/jxb/article/64/8/2423/645638.

21. Matton, D. P. et al. Hypervariable Domains of Self-Incompatibility RNases Mediate Allele-Specific Pollen Recognition. The Plant Cell Online 9, 1757–1766 (1997). URL http://www.plantcell.org/content/9/10/1757.

22. Matton, D. P. et al. Production of an S RNase with Dual Specificity Suggests a Novel Hypothesis for the Generation of New S Alleles. The Plant Cell 11, 2087–2097 (1999). URL http://www.plantcell.org/content/11/11/2087.

23. Bodčvá, K., Priklopil, T., Field, D. L., Barton, N. H. & Pickup, M. Evolutionary Pathways for the Generation of New Self-Incompatibility Haplotypes in a Non-self Recognition System. Genetics genetics.300748.2018 (2018). URL http://www.genetics.org/content/early/2018/04/30/genetics.118.300748.

24. Harkness, A., Goldberg, E. E. & Brandvain, Y. Diversification or Collapse of Self-Incompatibility Haplotypes as a Rescue Process. The American Naturalist 197, E89–E109 (2020). URL https://www.journals.uchicago.edu/doi/full/10.1086/712424. Publisher: The University of Chicago Press.

25. Fontana, W. & Schuster, P. Shaping Space: the Possible and the Attainable in RNA Genotype–phenotype Mapping. Journal of Theoretical Biology 194, 491–515 (1998). URL https://www.sciencedirect.com/science/article/pii/S0022519398907718.

26. Ancel, L. W., Fontana, W. et al. Plasticity, evolvability, and modularity in RNA. Journal of Experimental Zoology 288, 242–283 (2000).

27. Wagner, A. Robustness and evolvability: a paradox resolved. Proceedings of the Royal Society B: Biological Sciences 275, 91–100 (2008). URL http://rspb.royalsocietypublishing.org/content/275/1630/91.short.

28. Lancet, D., Sadovsky, E. & Seidemann, E. Probability Model for Molecular Recognition in Biological Receptor Repertoires: Significance to the Olfactory System. Proceedings of the National Academy of Sciences 90, 3715–3719 (1993). URL http://www.pnas.org/content/90/8/3715.

29. Lässig, M. From biophysics to evolutionary genetics: statistical aspects of gene regulation. BMC Bioinformatics 8, 1–21 (2007). URL 10.1186/1471-2105-8-S6-S7.

30. Friedlander, T., Prizak, R., Barton, N. H. & Tkačik, G. Evolution of new regulatory functions on biophysically realistic fitness landscapes. Nature Communications 8, 216 (2017). URL https://www.nature.com/articles/s41467-017-00238-8.

31. Wainwright, P. C., Alfaro, M. E., Bolnick, D. I. & Hulsey, C. D. Many-to-one mapping of form to function: A general principle in organismal design?1. Integrative and Comparative Biology 45, 256–262 (2005). URL 10.1093/icb/45.2.256. https://academic.oup.com/icb/article-pdf/45/2/256/2336200/i1540-7063-045-02-0256.pdf.

32. Elhanati, Y., Murugan, A., Callan, C. G., Mora, T. & Walczak, A. M. Quantifying selection in immune receptor repertoires. Proceedings of the National Academy of Sciences 111, 9875– 9880 (2014). URL http://www.pnas.org/content/111/27/9875.

33. Erez, K., Jangid, A., Feldheim, O. N. & Friedlander, T. The role of promiscuous molecular recognition in the evolution of RNase-based self-incompatibility in plants. Nature Communications 15, 4864 (2024). URL https://www.nature.com/articles/s41467-024-49163-7. Publisher: Nature Publishing Group.

34. Cover, T. M. & Thomas, J. A. Elements of information theory (John Wiley & Sons, 2012).

35. Dembo, A. & Zeitouni, O. Large Deviations Techniques and Applications, vol. 38 of Stochastic Modelling and Applied Probability (Springer, Berlin, Heidelberg, 2010). URL http://link.springer.com/10.1007/978-3-642-03311-7.

36. Berg, J., Willmann, S. & Lässig, M. Adaptive evolution of transcription factor binding sites. BMC Evolutionary Biology 4, 42 (2004). URL 10.1186/1471-2148-4-42.

37. Sella, G. & Hirsh, A. E. The application of statistical physics to evolutionary biology. Proceedings of the National Academy of Sciences of the United States of America 102, 9541– 9546 (2005). URL http://www.pnas.org/content/102/27/9541.

38. Zhao, H. et al. Origin, loss, and regain of self-incompatibility in angiosperms. The Plant Cell 34, 579–596 (2022). URL 10.1093/plcell/koab266.

39. Lukatsky, D. B., Shakhnovich, B. E., Mintseris, J. & Shakhnovich, E. I. Structural Similarity Enhances Interaction Propensity of Proteins. Journal of Molecular Biology 365, 1596–1606 (2007). URL https://www.sciencedirect.com/science/article/pii/S002228360601549X.

40. Peleg, O.Choi, J.-M. & Shakhnovich, E. I. Evolution of Specificity in Protein-Protein Interactions. Biophysical Journal 107, 1686–1696 (2014). URL https://www.sciencedirect.com/science/article/pii/S0006349514008418.

41. Deeds, E. J. & Rowland, M. A. The Evolution of Crosstalk in Signaling Networks. Biophysical Journal 106, 643a (2014). URL http://www.cell.com/biophysj/abstract/S0006-3495(13)04819-4.

42. Aakre, C. et al. Evolving New Protein-Protein Interaction Specificity through Promiscuous Intermediates. Cell 163, 594–606 (2015). URL http://www.sciencedirect.com/science/article/pii/S0092867415012726.

43. Lt uksza, M. & Lässig, M. A predictive fitness model for influenza. Nature 507, 57–61 (2014). URL https://www.nature.com/articles/nature13087.

44. Holding, M. L., Drabeck, D. H., Jansa, S. A. & Gibbs, H. L. Venom Resistance as a Model for Understanding the Molecular Basis of Complex Coevolutionary Adaptations. Integrative and Comparative Biology 56, 1032–1043 (2016). URL https://academic.oup.com/icb/article/56/5/1032/2420622. Publisher: Oxford Academic.

45. Arbuckle, K., Rodríguez de la Vega, R.C. & Casewell, N. R. Coevolution takes the sting out of it: Evolutionary biology and mechanisms of toxin resistance in animals. Toxicon 140, 118–131 (2017). URL http://www.sciencedirect.com/science/article/pii/S0041010117303367.

46. Nguyen, C. C. & Saier, M. H. Phylogenetic, structural and functional analyses of the LacI-GalR family of bacterial transcription factors. FEBS Letters 377, 98–102 (1995). URL http://onlinelibrary.wiley.com/doi/10.1016/0014-5793(95)01344-X/abstract.

47. Sear, R. P. Highly specific protein–protein interactions, evolution and negative design. Physical Biology 1, 166 (2004). URL http://stacks.iop.org/1478-3975/1/i=3/a=004.

48. Sear, R. P. Specific protein–protein binding in many-component mixtures of proteins. Physical Biology 1, 53 (2004). URL http://stacks.iop.org/1478-3975/1/i=2/a=001.

49. George, J. T., Kessler, D. A. & Levine, H. Effects of thymic selection on T cell recognition of foreign and tumor antigenic peptides. Proceedings of the National Academy of Sciences 114, E7875–E7881 (2017). URL http://www.pnas.org/content/114/38/E7875.

50. Friedlander, T., Prizak, R., Guet, C. C., Barton, N. H. & Tkačik, G. Intrinsic limits to gene regulation by global crosstalk. Nature Communications 7, 12307 (2016). URL http://www.nature.com/ncomms/2016/160804/ncomms12307/abs/ncomms12307.html.

51. McClune, C. J. & Laub, M. T. Constraints on the expansion of paralogous protein families. Current Biology 30, R460–R464 (2020). URL https://www.cell.com/current-biology/abstract/S0960-9822(20)30282-7. Publisher: Elsevier.

52. Maerkl, S. J. & Quake, S. R. A Systems Approach to Measuring the Binding Energy Landscapes of Transcription Factors. Science 315, 233–237 (2007). URL http://www.sciencemag.org/cgi/content/abstract/sci;315/5809/233.

53. Pougach, K. et al. Duplication of a promiscuous transcription factor drives the emergence of a new regulatory network. Nature Communications 5, 4868 (2014). URL http://www.nature.com/ncomms/2014/140910/ncomms5868/full/ncomms5868.html?WT.ec_id=NCOMMS-20140917.

54. Shultzaberger, R. K., Maerkl, S. J., Kirsch, J. F. & Eisen, M. B. Probing the Informational and regulatory plasticity of a transcription factor DNA–binding domain. PLoS genetics 8 (2012). URL http://europepmc.org/articles/PMC3315485.

55. Adams, R. M., Mora, T., Walczak, A. M. & Kinney, J. B. Measuring the sequence-affinity landscape of antibodies with massively parallel titration curves. eLife 5, e23156 (2016). URL https://elifesciences.org/content/5/e23156v3.

56. Mayer, A., Balasubramanian, V., Mora, T. & Walczak, A. M. How a well-adapted immune system is organized. Proceedings of the National Academy of Sciences 112, 5950–5955 (2015). URL https://www.pnas.org/doi/full/10.1073/pnas.1421827112. Publisher: Proceedings of the National Academy of Sciences.

57. Noivirt-Brik, O., Unger, R. & Horovitz, A. Analysing the origin of long-range interactions in proteins using lattice models. BMC Structural Biology 9, 4 (2009). URL 10.1186/1472-6807-9-4.

