## supplementary information for "Reconciling conflicting selection pressures in the plant collaborative non-self recognition self-incompatibility system"

#### Supplementary information file contains the following,

##### Supplementary notes

1. Classification of the 20 amino acids into four biochemical classes.
2. The probability that offspring are formed by self-fertilized self-compatible, out-crossed self-compatible, and self-incompatible female S-haplotypes.
3. Haploid vs. diploid models.
4. Empirical genomic data.

##### Supplementary figures

Fig. S1: Within-run and between-run quantitative measures are very close, suggesting ergodicity of the system.

Fig. S2: The interaction energy threshold ($E_{\mathrm{th}}$) biases the frequencies of RNases amino acids whereas the frequencies of SLFs amino acids remain almost neutral.

Fig. S3: The frequency of hydrophobic AA in RNases and SLFs against the interaction energy threshold $E_{\mathrm{th}}$ in the reduced model behaves very similarly to the full model.

Fig. S4: RNase AA frequencies in the reduced model with one SLF per S-haplotype are well-approximated by the analytical result.

Fig. S5: In the full model too, the frequency of hydrophobic AA $\psi_H$ in RNases decreases as the number of SLFs per S-haplotype increases, but their frequency in SLFs is insensitive to the number of SLFs per S-haplotype.

Fig. S6: The amino acid frequencies are insensitive to the inbreeding depression $\delta$ in a broad range.

Fig. S7: Local arrangement in hydrophobic and neutrally polarized amino acids is observed amongst partner and same S-haplotype proteins.

Fig. S8: The number of fertilization attempts per female affects the distributions of interaction energies and the RNase (but not the SLF) amino acid composition

Fig. S9: Within S-haplotype energy distribution amongst sequences evolved in the simulation under the reduced model compared to the neutral case.

Fig. S10: The number of distinct RNases detoxified by a single SLF increases with promiscuity.

Fig. S11: The population proportion of unclassified S-haplotypes decreases, and the proportion of self-compatible ones increases with $E_{\mathrm{th}}$.

Fig. S12: The divergence of the RNase AA frequencies reduced relative to the prior increase with the number of SLF genes per S-haplotype and witn $E_{\mathrm{th}}$ in both the full and the reduced models.

Fig. S13: Compatible fraction amongst RNase-SLF pairs in different scenarios.

Table S1: Classification of the 20 amino acids into four biochemical classes.

| bio-classes | H (%) | P (%) | + (%) | - (%) |
|---|---|---|---|---|
| amino | F (3.86) | C (1.37) | E (6.74) | R (5.53) |
| acids | W (1.09) | Y (2.92) | D (5.46) | K (5.82) |
| | M (2.41) | T (5.35) | | |
| | L (9.65) | S (6.60) | | |
| | I (5.93) | N (4.06) | | |
| | V (6.86) | H (2.27) | | |
| | G (7.08) | Q (3.93) | | |
| | P (4.72) | | | |
| | A (8.26) | | | |

#### The probability that offspring are formed by either self-fertilized self-compatible, out-crossed self-compatible, and self-incompatible female S-haplotypes

Consider $x$ and $y$ to be the population fractions of SC and SI S-haplotypes at a given generation where $x + y = 1$. We calculate here the proportions of offspring produced by either self-fertilized self-compatible female $f^{\rm s}_{\rm SC}$, out-crossed self-compatible female $f^{\rm ns}_{\rm SC}$, and out-crossed self-incompatible female $f_{\rm SI}$:

$$\begin{aligned} f^{\rm s}_{\rm SC} &= x\ \alpha\ (1-\delta) + \overbrace{\underbrace{(x\ \alpha\ \delta)}_{\text{dying fraction}}\ \underbrace{x\ \alpha\ (1-\delta)}_{\text{self-fertilized}}}^{\text{round 1}} \\ &\quad + \overbrace{\underbrace{(x\ \alpha\ \delta)^2}_{\text{dying fraction}}\ \underbrace{x\ \alpha\ (1-\delta)}_{\text{self-fertilized}}}^{\text{round 2}} + \cdots \end{aligned} \tag{1}$$

$$\begin{aligned} f^{\rm ns}_{\rm SC} &= x\ (1-\alpha) + \overbrace{\underbrace{(x\ \alpha\ \delta)}_{\text{dying fraction}}\ \underbrace{x\ (1-\alpha)}_{\text{out-crossed}}}^{\text{round 1}} \\ &\quad + \overbrace{\underbrace{(x\ \alpha\ \delta)^2}_{\text{dying fraction}}\ \underbrace{x\ (1-\alpha)}_{\text{out-crossed}}}^{\text{round 2}} + \cdots \end{aligned} \tag{2}$$

$$\begin{aligned} f_{\rm SI} &= y + \overbrace{\underbrace{(x\ \alpha\ \delta)}_{\text{dying fraction}}\ \underbrace{y}_{\text{self-incompatible}}}^{\text{round 1}} \\ &\quad + \overbrace{\underbrace{(x\ \alpha\ \delta)^2}_{\text{dying fraction}}\ \underbrace{y}_{\text{self-incompatible}}}^{\text{round 2}} + \cdots \end{aligned} \tag{3}$$

The first term in the right-hand side of the equations is the fraction of $f^{\rm s}_{\rm SC}$, $f^{\rm ns}_{\rm SC}$, and $f_{\rm SI}$. The second term represents the fraction of $f^{\rm s}_{\rm SC}$, $f^{\rm ns}_{\rm SC}$, and $f_{\rm SI}$ S-haplotypes being chosen after $x\ \alpha\ \delta$ fraction of SC S-haplotypes died - could not make up to adulthood due to inbreeding depression. The third term is the same as the second but the fraction of S-haplotypes that could not make up to adulthood is $(x\ \alpha\ \delta)^2$. Eqs. (1), (2) and (3) lead to,

$$f^{\rm s}_{\rm SC} = x\ \alpha\ (1-\delta)\ [1 + (x\ \alpha\ \delta) + (x\ \alpha\ \delta)^2 + \cdots] \tag{4}$$

$$f^{\rm ns}_{\rm SC} = x\ (1-\alpha)\ [1 + (x\ \alpha\ \delta) + (x\ \alpha\ \delta)^2 + \cdots] \tag{5}$$

$$f_{\rm SI} = y\ [1 + (x\ \alpha\ \delta) + (x\ \alpha\ \delta)^2 + \cdots] \tag{6}$$

The probability of choosing the offspring maternal parent to be self-fertilized self-compatible $f^{\rm s}_{\rm SC}$, out-crossed self-compatible $f^{\rm ns}_{\rm SC}$, and self-incompatible $f_{\rm SI}$ S-haplotype are given as,

$$f^{\rm s}_{\rm SC} = \frac{x\ \alpha\ (1-\delta)}{1 - x\ \alpha\ \delta} \tag{7}$$

$$f^{\rm ns}_{\rm SC} = \frac{x\ (1-\alpha)}{1 - x\ \alpha\ \delta} \tag{8}$$

$$f_{\rm SI} = \frac{1-x}{1 - x\ \alpha\ \delta} \tag{9}$$

##### Haploid vs. diploid models

Table S2: Comparison between the haploid model of this paper and the diploid model of Erez et al. The models differ in the definition of an individual as either a diploid or a haploid, but their qualitative behaviors are very similar.

| **Model construction** | | |
|---|---|---|
| **Factor** | **Diploid model** | **Haploid model** |
| An individual in the population consists of | two S-haplotypes | one S-haplotype |
| To fertilize a maternal plant a paternal S-haplotype should detoxify | two RNases (of both S-haplotypes) | one RNase |
| Duplication and deletion of SLF per S-haplotype | enabled | disabled |
| The number of SLFs per S-haplotype | variable (if gene duplication and deletion are enabled) | fixed |
| **Results:** | | |
| The vast majority of the population self-organizes into genetically heterogeneous compatibility classes | applicable; ≈ 7 classes for $2N$ = 1000 S-haplotypes at $E_{\text{th}}$ = -6 | applicable; ≈ 6 classes for $N$ = 1000 S-haplotypes at $E_{\text{th}}$ = -6 |
| The number of classes increases with $E_{\text{th}}$ | applicable; | applicable; |
| The fraction of unclassified S-haplotypes | ≈ 5% for $2N$ = 1000 S-haplotypes at $E_{\text{th}}$ = -6 | ≈ 6.8% for $N$ = 1000 S-haplotypes at $E_{\text{th}}$ = -6 |
| Average effective number of distinct RNases | ≈ 8 RNases for $2N$ = 1000 S-haplotypes at $E_{\text{th}}$ = -6 | ≈ 7 RNases for $N$ = 1000 S-haplotypes at $E_{\text{th}}$ = -6 |

#### Genomic data

Table S3: We used in our analysis the RNase amino acids reported by Vieira *et al.* [1] to be positively selected, and classified them into either of the four biochemical classes following Table S1.

| **Data set 1** | **Sequence** |
|---|---|
| $S3$-*RNase P. integrifolia* | EHFDD-KVSDVRFDKFKIKTSNLRK |
| $S7$-*RNase P. integrifolia* | VSTKKPKITGLKYKPYRNERQKQIN |
| $S12$-*RNase P. integrifolia* | EHYTK-KAPDVRFDHYKINNSNLRK |
| $S13$-*RNase P. integrifolia* | KHFTD-KTRDINFDKYKDKGKSLRN |
| $S2$-*RNase P. integrifolia* | KHFTD-KSRDIRFDKYSTKEKNLEN |
| $S11$-*RNase P. integrifolia* | QHVAK-TQKVIDKSSETRTQTERQK |
| **Data set 2** | **Sequence** |
| $S5$,$SB1$-*RNase P. hybrida* | YSTTDEVKDKKTDADKGTKLSSEK |
| $S11$-*RNase P. hybrida* | QHVA-TQKVIDKSSETRTQTERQK |
| $S17$-*RNase P. axillaris* | ERQADYYVGKHFNEVLKYHSNLTN |
| $S7$-*RNase P. hybrida* | QHVD-TTTAKAYTRDIQLRSEREK |
| **Data set 3** | **Sequence** |
| $S10$-*RNase P. hybrida* | EQRT-TESLGLRFDETMQDHENRMN |
| $S19$-*RNase P. axillaris* | VKTDSTKNKAKKSLHILTSKEPAST |
| $S7$-*RNase P. hybrida* | QHVD–TTTAKAYTRDIQLRSEREK |
| $S11$-*RNase P. hybrida* | QHVA–TQKVIDKSSETRTQTERQK |
| $S17$-*RNase P. axillaris* | ERQADYKVNGKHFNEVLKYHSNLTN |

### Supplementary figures

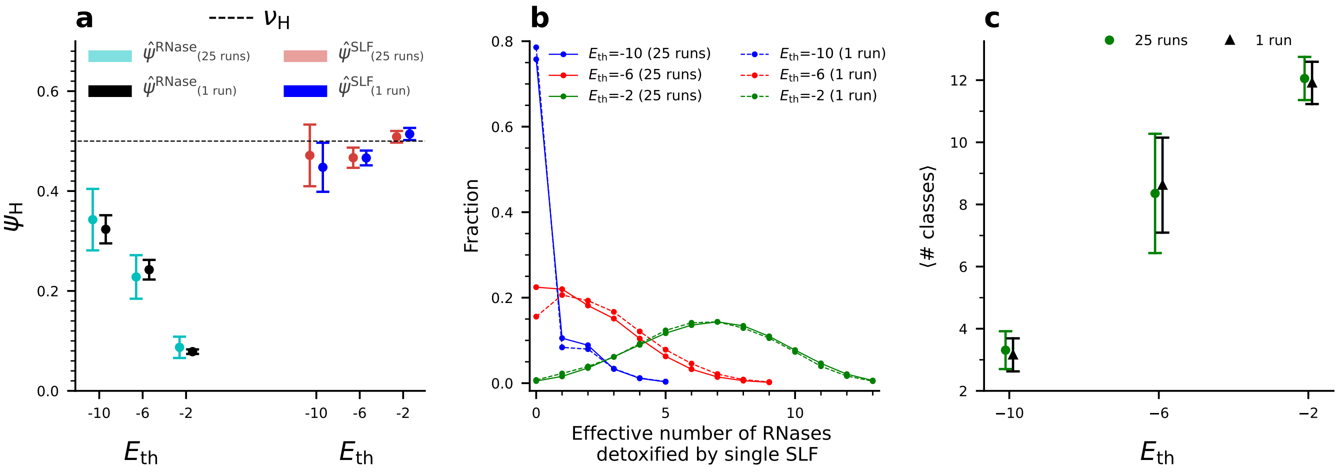


Figure S1: **Within-run and between-run quantitative measures are very close, suggesting ergodicity of the system.** We compare different quantitative measures used in this paper calculated either using a single run data or 25 independent runs lumped together. **(a)** The frequency of hydrophobic amino acids, **(b)** the distribution of the effective number of RNase partners per SLF, and **(c)** the average number of classes for 25 and 1 runs for three different energy threshold values. In all measures we observe that the distribution of a single run well-approximates the distribution over 25 distinct runs, suggesting ergodicity of the system. The distributions are based on 25 independent runs for each $E_{\mathrm{th}}$ value, 1200 data points taken from each run, with a 25-generation interval between consecutive time points used for the analysis. The single-run results were obtained by continuing one of the simulations from the time-point where it was stopped for additional $10^5$ generations. We plotted the figure using data 25-generation intervals. Parameter values used are the default values listed in Table 2 main text.

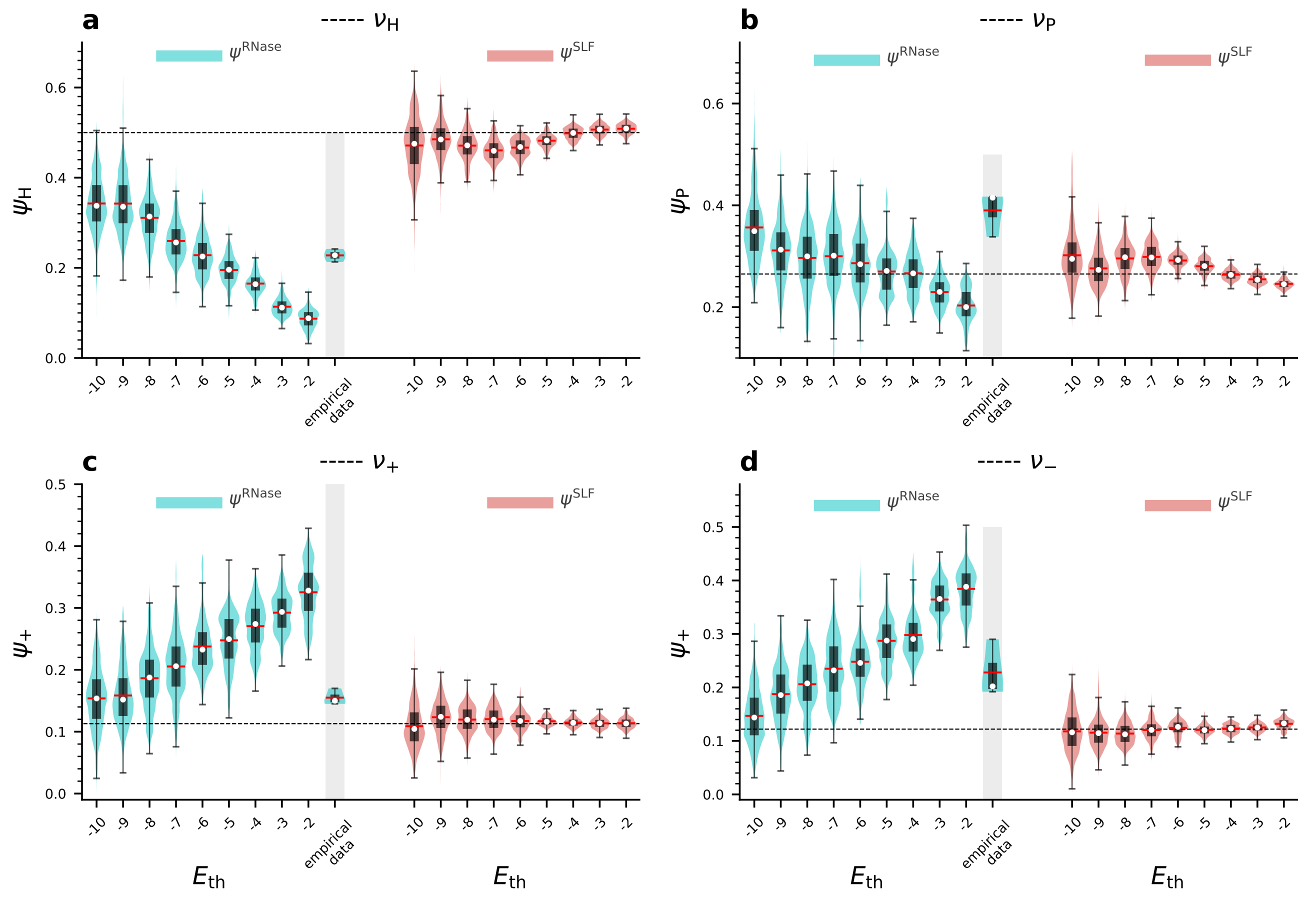


Figure S2: **The interaction energy threshold ($E_{th}$) biases the frequencies of RNases amino acids whereas the frequencies of SLFs amino acids remain almost neutral**. We plot here the frequencies of hydrophobic (a), neutral polar (b), positively charged (c), and negatively charged (d) amino acids of RNases (cyan, panel left side) and functional SLFs (red, panel right side) against $E_{th}$ obtained in populations evolved under the full model. The red line and white circle in the violin plot refer to the distribution median and mean, respectively. The dashed black lines correspond to the prior frequencies of the relevant amino acid categories. In the RNases, we observe an increase in hydrophobic and a decrease in charged amino acids with the increase in $E_{th}$. In contrast, in the SLFs, all four AA categories SLFs are almost insensitive on average to $E_{th}$. For the RNases, we also show the empirical amino acid frequencies (grey background) as determined in several studies (see Methods for details). The distributions are based on 25 independent runs for each $E_{th}$ value, 1200 data points taken from each run, with a 25-generation interval between consecutive time points used for the analysis. All these data points were taken after the entire population descends from a single common ancestor and the amino acid frequencies have already reached their steady-state values. The total runtime for each is $2 \times 10^5$ generations. Parameter values are as in Table 2 main text.

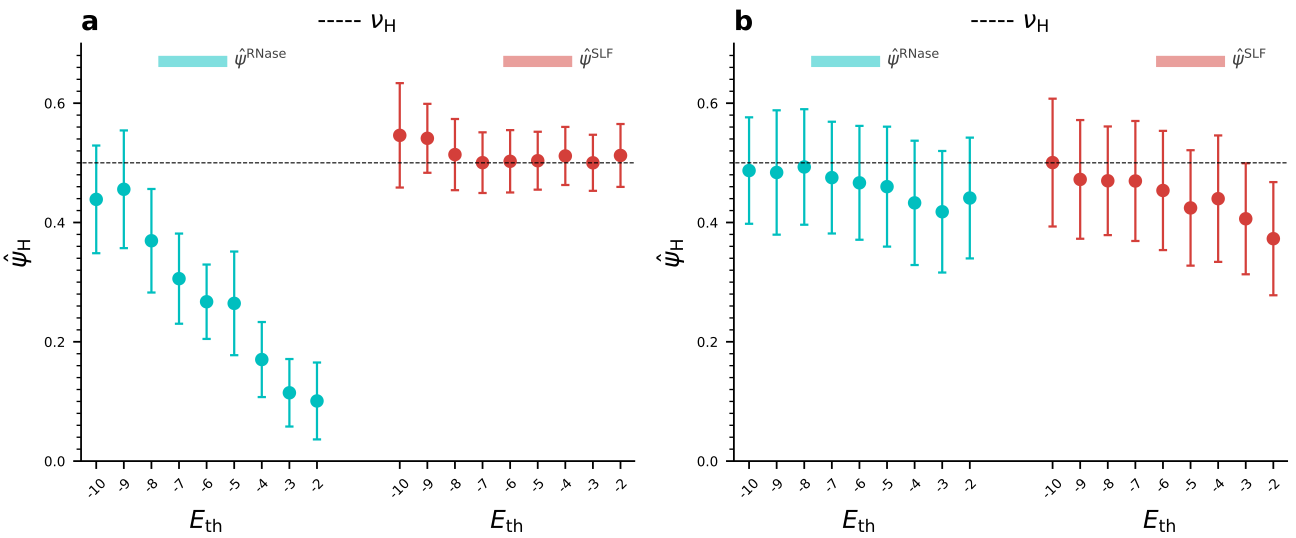


Figure S3: **The frequency of hydrophobic AA in RNases and SLFs against the interaction energy threshold $E_{\mathrm{th}}$ in the reduced model behaves very similarly to the full model**. The frequency of hydrophobic AA in RNases (cyan) and SLFs (red) in populations evolved in the reduced model (only selection against self-compatibility but no reproductive cycle) against $E_{\mathrm{th}}$ with either 9 **(a)** or 1 **(b)** SLFs per S-haplotype. As $E_{\mathrm{th}}$ becomes higher the pressure to avoid self-compatibility strengthens, and consequently the frequency of hydrophobic amino acids decreases in the RNases, but not in the SLFs. This dependence on $E_{\mathrm{th}}$ and asymmetry between RNase and SLF is very similar to the behavior of the full model as shown in Fig. S2a. The horizontal dashed line marks the prior frequency of hydrophobic AA, $\nu_H$. Circles denote averages, bars represent ±STD relative to the mean. (a) is based on 25 independent runs for each number of SLFs per S-haplotype, with 1200 data points taken from each run, with a 25-generation interval between consecutive time points used for the analysis. (b) is based on 200 independent runs for each number of SLFs per S-haplotype, with 200 data points taken from each run, with a 250-generation interval between consecutive time points used for the analysis. All these data points were taken after the entire population descends from a single common ancestor and the amino acid frequencies have already reached their steady-state values. The total runtime is $1.5 \times 10^5$, and $2.0 \times 10^5$ generations for (a), and (b) respectively. Parameter values are as in Table 2 main text.

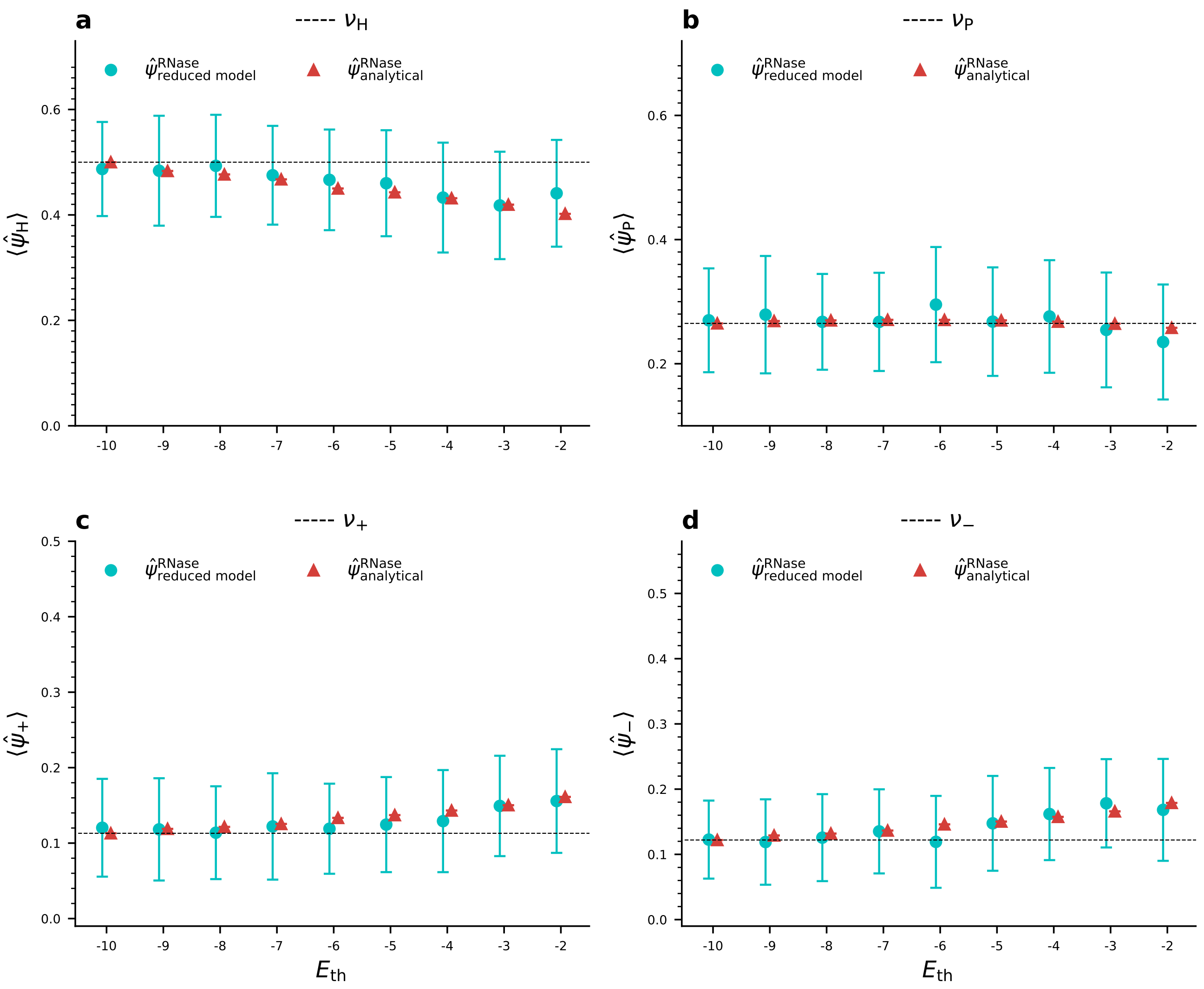


Figure S4: **RNase AA frequencies in the reduced model with one SLF per S-haplotype are well-approximated by the analytical result**. We plot the AA frequencies with one SLF per S-haplotype obtained in simulations of the reduced model (cyan circle) and in the analytical approximation (red triangle). The horizontal dashed lines mark the prior frequencies of the corresponding AAs. Circles and triangles denote averages in the corresponding model, bars represent ±STD relative to the mean. The figure is based on 25 independent runs for each number of SLFs per S-haplotype, with 450 data points taken from each run, with a 25-generation interval between consecutive time points. All these data points were taken after the entire population descends from a single common ancestor and the amino acid frequencies have already reached their steady-state values. The total runtime for each $E_{\rm th}$ value is $1.5 \times 10^5$ generations. Parameter values are as in Table 2 main text.

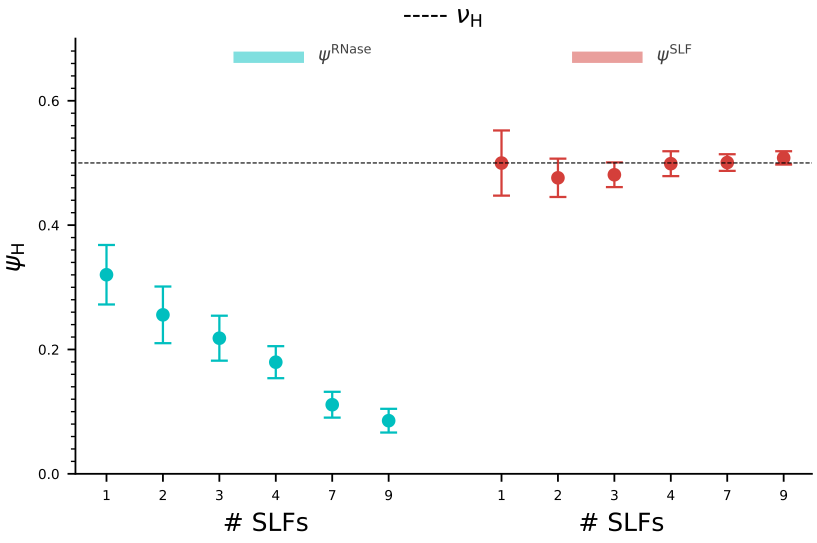


Figure S5: **In the full model too, the frequency of hydrophobic AA $\psi_H$ in RNases decreases as the number of SLFs per S-haplotype increases, but their frequency in SLFs is insensitive to the number of SLFs per S-haplotype**. The frequency of hydrophobic AA in RNases (cyan) and SLFs (red) in evolved populations against the number of SLFs per S-haplotype. As the number of SLFs per S-haplotype increases the pressure to avoid self-compatibility strengthens, and consequently the frequency of hydrophobic amino acids decreases. The horizontal dashed line marks the prior frequency of hydrophobic AAs. Circles denote averages, bars represent $\pm$STD relative to the mean. The figure is based on 16 independent runs for each number of SLFs per S-haplotype, with 1200 data points taken from each run, with a 25-generation interval between consecutive time points used for the analysis. All these data points were taken after the entire population descends from a single common ancestor and the amino acid frequencies have already reached their steady-state values. The total runtime for each is $4.5 \times 10^5$ generations. Parameter values are as in Table 2 main text.

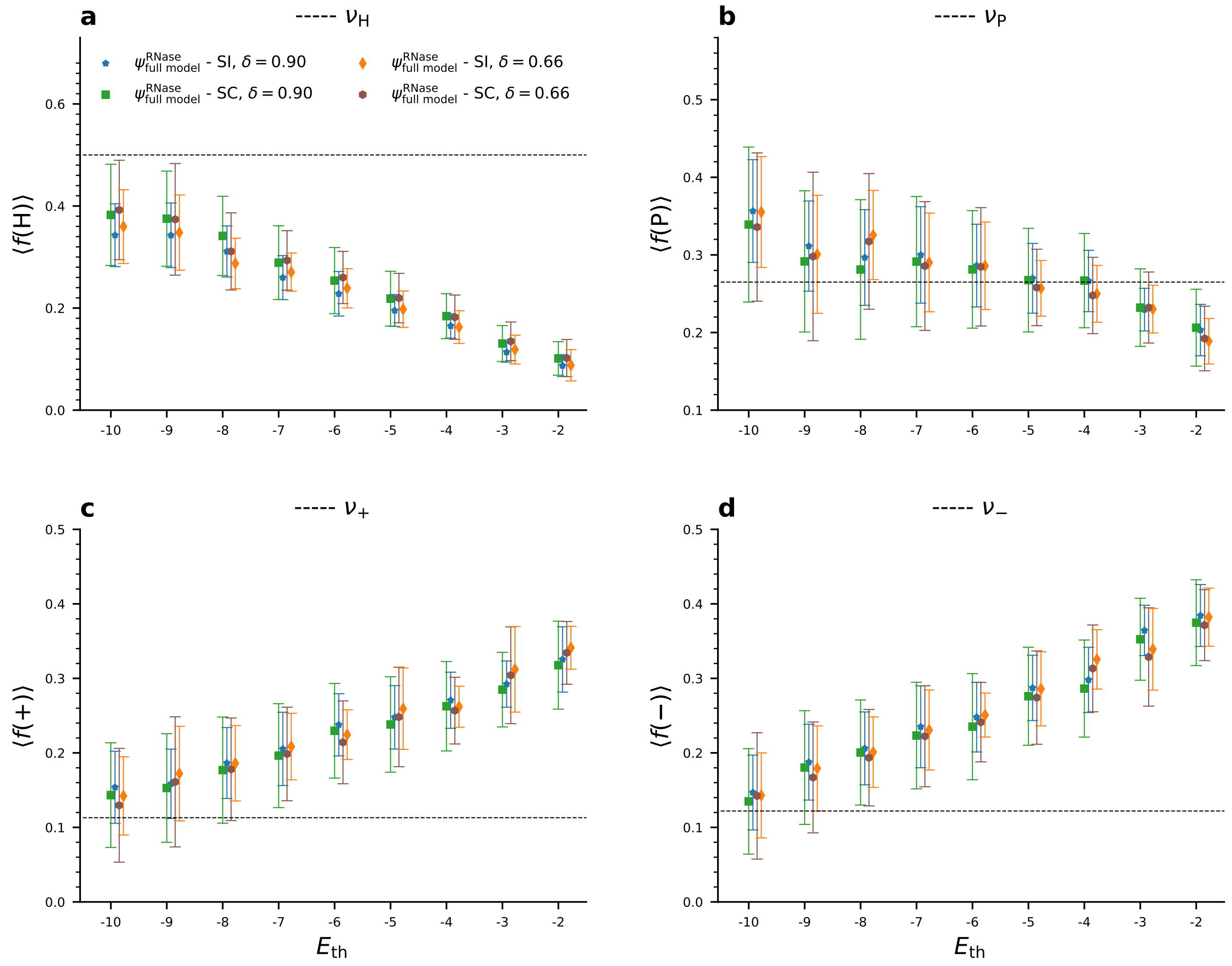


Figure S6: **The amino acid frequencies are insensitive to the inbreeding depression $\delta$ in a broad range**. The AA frequencies obtained in simulations of the full model under two different inbreeding depression values $\delta = 0.90$, and 0.66 for self-incompatible and self-compatible S-haplotypes. We find that for the self-incompatible S-haplotypes the frequencies under the two different $\delta$ values are very close. We also observe that the AA frequencies of the self-compatible S-haplotypes are quite close to those of the self-incompatible ones, suggesting perpetual transitions between the SI and SC sub-populations, such that their AAS frequencies are similar. The horizontal dashed lines mark the prior frequencies, $\nu$. Different markers denote averages, bars represent ±STD relative to the mean. In the figure, data for $\delta = 0.90$ (blue star and green square) are based on 25 independent runs, and data for $\delta = 0.66$ (orange diamond and brown hexagon) are based on 10 independent runs, with 1000 data points taken from each run, with a 25-generation interval between consecutive time points used for the analysis. All these data points were taken after the entire population descended from a single common ancestor and the amino acid frequencies had already reached their steady-state values. The total runtime for each is $E_{th}$ is the same as in Table 4. Parameter values are as in Table 2 main text.

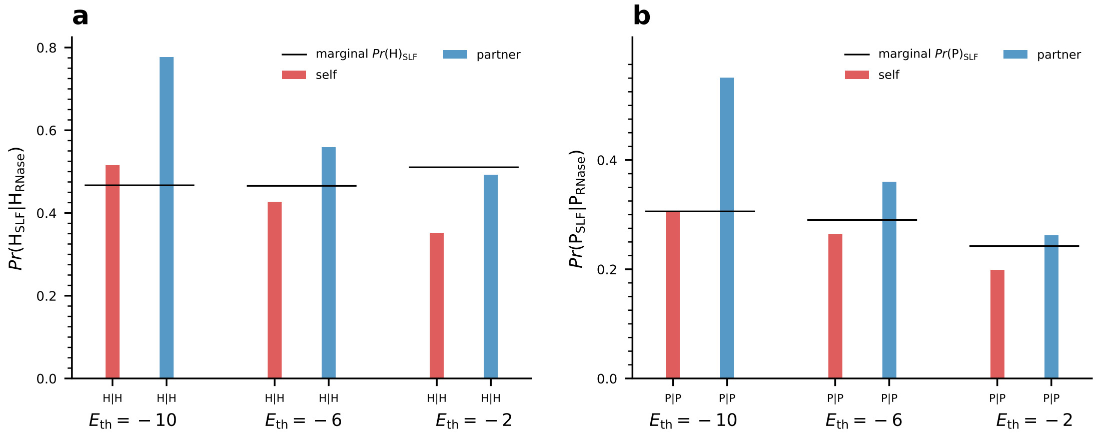


Figure S7: **Local arrangement of hydrophobic and neutrally polarized amino acids is observed amongst partner and same S-haplotype proteins.** **(a)** Frequencies of hydrophobic (H) amino acids in SLFs conditioned on RNase having hydrophobic amino in the corresponding position for protein pairs of either for partners from different S-haplotypes for various $E_{\mathrm{th}}$ values. The horizontal lines mark the marginal H frequencies in SLFs. **(b)** Similar to (a) but for neutrally polarized amino acids P. Both H-H and P-P pairs promoter attraction, and indeed we find their enrichment for partner protein pairs, mostly under low $E_{\mathrm{th}}$ where the selection to match dominates. Complementary to that, we find their under-representation amongst same S-haplotype RNase-SLF pairs, mostly under high $E_{\mathrm{th}}$, where selection to avoid self-compatibility dominates. These results demonstrate the use of the local strategy, which is most pronounced for partner attraction under low promiscuity, complementary to Fig. 5, where its effect on the charged AA arrangement was illustrated. The calculations are based on 25 independent runs for each $E_{\mathrm{th}}$ value, with 600 data points from each run with 25-generation intervals between consecutive time points. All the data points used in the analysis are beyond the times in which the entire population descends from a single ancestor and the frequency of amino acid frequencies have reached their steady-state values. Run times are listed in Table 4 main text. We used the default parameter values listed in Table 2 main text, except for $E_{\mathrm{th}}$.

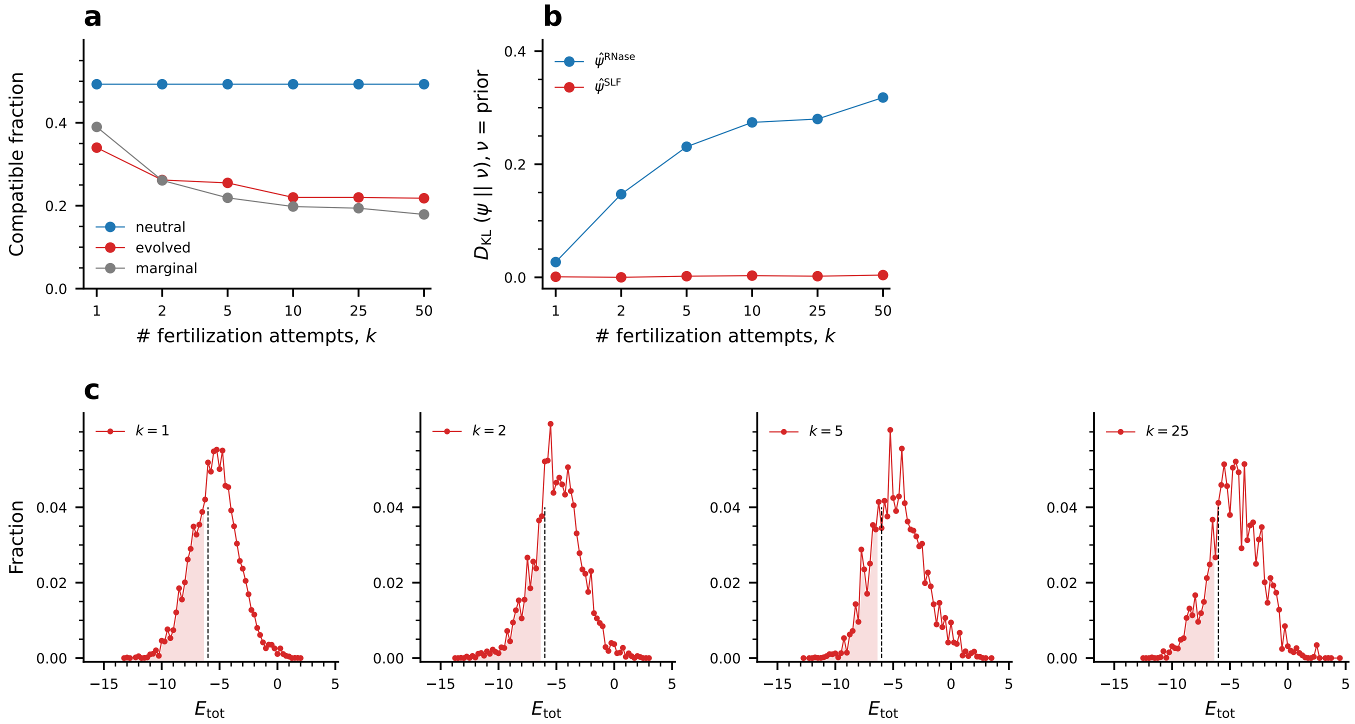


Figure S8: **The number of fertilization attempts per female affects the distributions of interaction energies and the RNase (but not the SLF) amino acid composition**. **(a)** The fraction of compatible SLF-RNase pairs amongst evolved sequences (red) and random sequences drawn from the marginal amino acid distributions of the evolved sequences (grey) for different numbers of fertilization attempts $k$. The compatible fraction amongst random sequences drawn from the prior distribution (blue) is shown for reference. We find that the compatible fraction decreases as fertilization attempts increase. This fraction saturates for $k \geq 10$. **(b)** The Kullback-Leibler divergences between the marginal AA distribution of evolved sequences and the prior AA distribution for RNases (blue) and functional SLFs (red) for different values of $k$. Similar to the results in the main text, the SLF AA distribution is also indistinguishable from the prior. In contrast, the RNase AA deviates from the prior, but this deviation is minor under pollen limitation ($k = 1$) and increases with $k$. **(c)** Distributions of interaction energies between evolved RNases and functional SLFs for all possible pairing combinations for different numbers of fertilization attempts $k$ per female. The shaded regions under the distributions mark the proportion of compatible pairings, namely $E_{\text{tot}} < E_{\text{th}}$, but only for the functional SLFs. The figure is based on 16 independent runs for each $k$ value, with 1200 data points taken from each run, with a 25-generation interval between consecutive time points used for the analysis. All these data points were taken after the entire population descends from a single common ancestor and the amino acid frequencies have already reached their steady-state values. The total runtime for each is $2 \times 10^5$ generations. Parameter values: interaction energy thresholds $E_{\text{th}} = -6$. The remaining parameters are as in Table 2 main text.

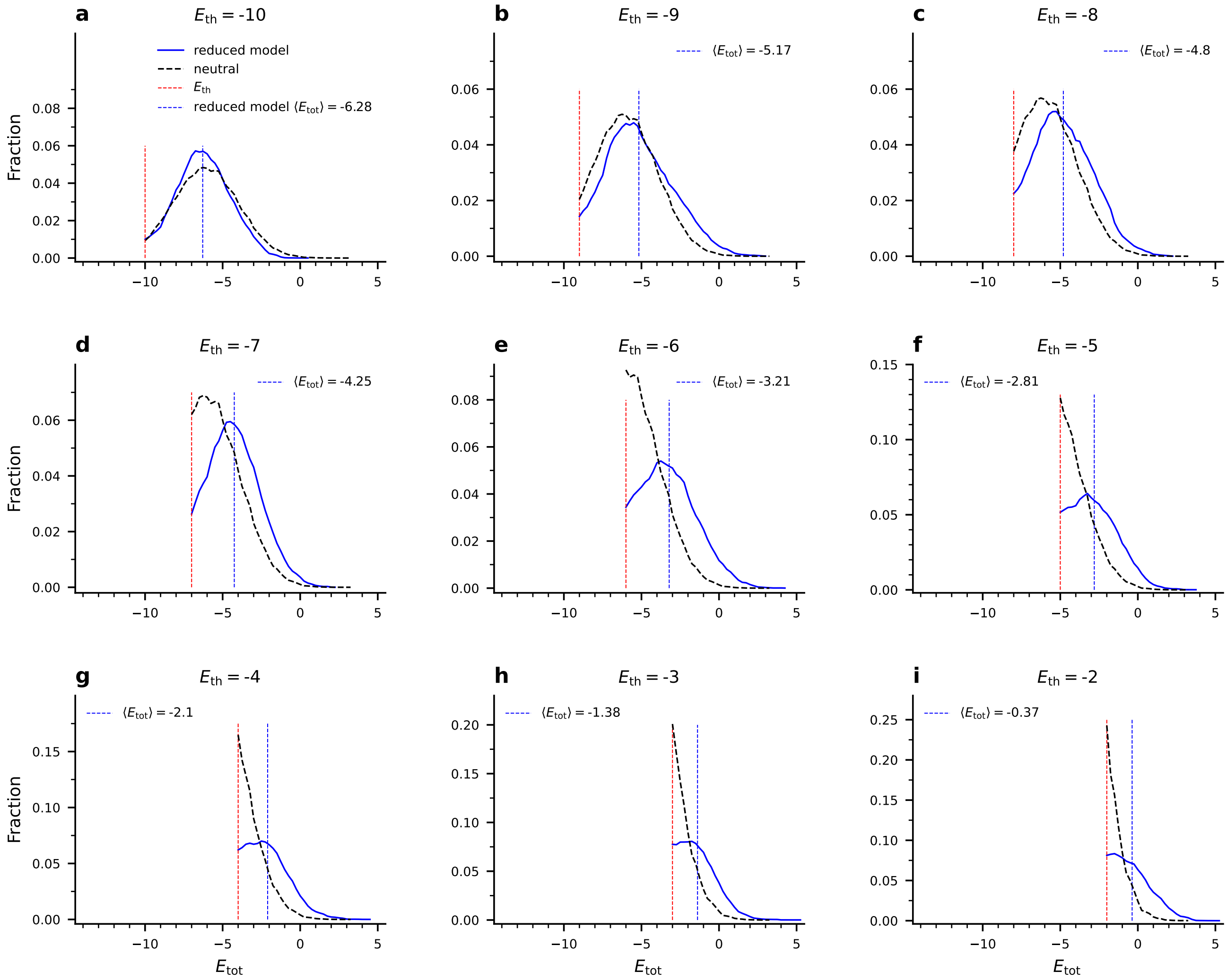


Figure S9: **Within S-haplotype energy distribution amongst sequences evolved in simulation under the reduced model compared to the neutral case**. For every $E_{\rm th}$ we plot the distribution of interaction energies $E_{\rm tot}$ between same-S-haplotype RNase and SLFs evolved in simulations under the reduced model, namely only under selection against self-compatibility (blue curves). The dashed vertical blue lines mark the distribution mean, and the red lines marks $E_{\rm th}$ in each case. The selection pressure constrains the interaction energies to exceed this value, hence the distributions are truncated at this value, as expected. We show for reference the energy distribution in the absence of any selection for $EE_{\rm tot} > E_{\rm th}$, scaled such that its integral equals 1 (dashed black curve). We observe that $E_{\rm tot}$ distribution evolved under low $E_{\rm th}$ is similar to the neutral one and the gap between $E_{\rm th}$ and the distribution mean is relatively large. As $E_{\rm th}$ increases the evolved distribution is driven away from the neutral one and the gap between $E_{\rm th}$ and the distribution mean decreases.

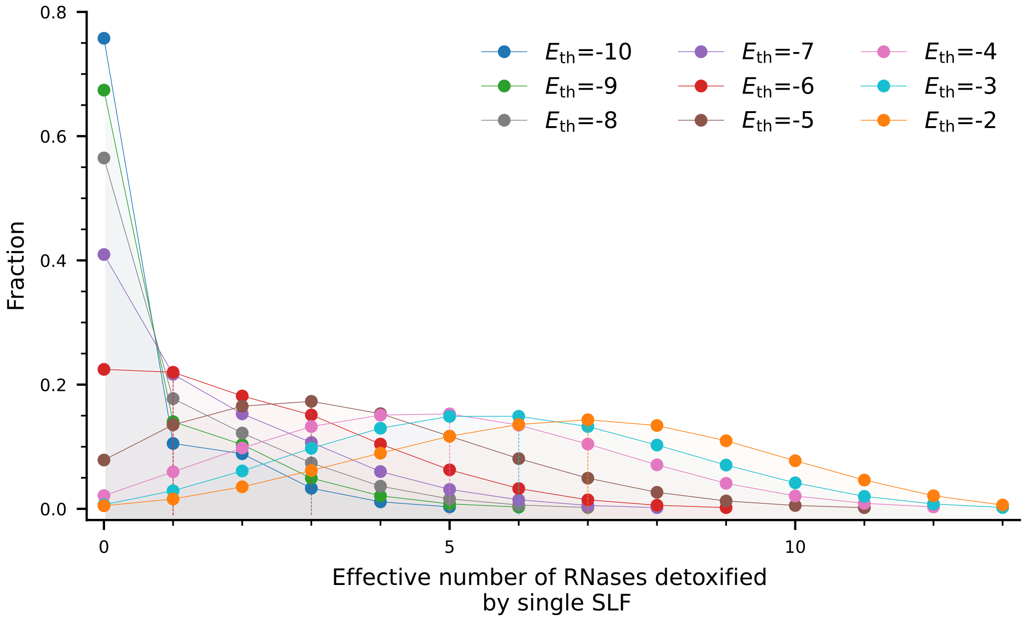


Figure S10: **The number of distinct RNases detoxified by a single SLF increases with promiscuity**. Distributions of the number of distinct RNases detoxified by a single SLF in the population for different values of $E_{\mathrm{th}}$, marked by different colors. The distributions are based on 25 independent runs for each $E_{\mathrm{th}}$ value, 1200 data points from each run with 25-generations interval between consecutive time points. The analysis only includes data points beyond the time that the entire population descended from a single ancestor and the amino acid frequencies have reached their final values. Run time is listed in Table 4 main text. Parameter values: the default values listed in Table 2 main text.

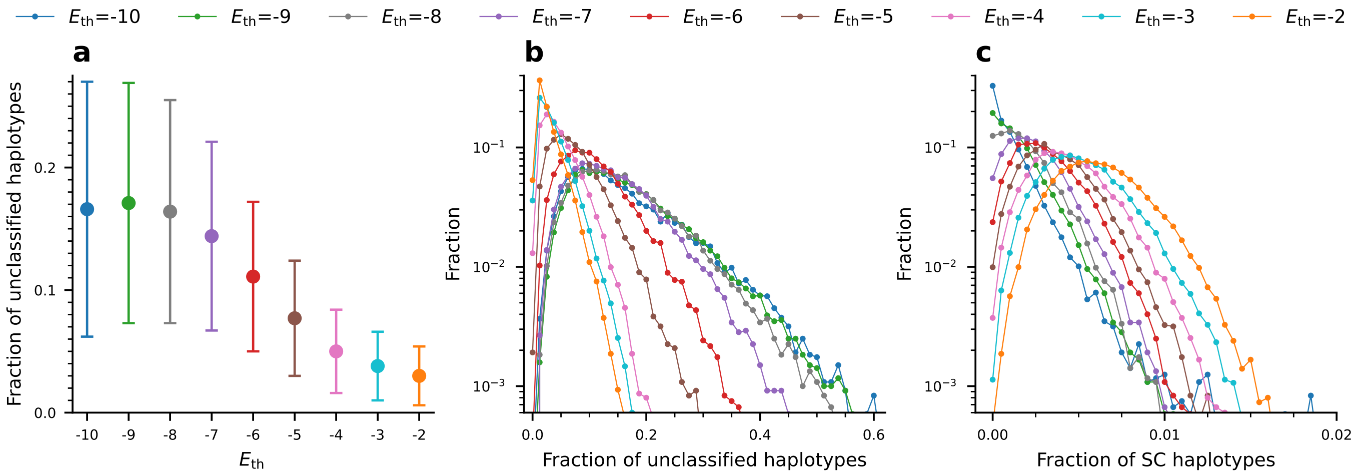


Figure S11: **The population proportion of unclassified S-haplotypes decreases, and the proportion of self-compatible ones increases with $E_{\text{th}}$. (a)** The average population proportion of unclassified S-haplotypes (either SI or SC) for different values of $E_{\text{th}}$, denoted by different colors. The colored circle and bar represent the mean and the ± STD. **(b-c)** Distributions of the population proportions of unclassified S-haplotypes (either SI or SC) (b) and self-compatible S-haplotypes (c) for different values of $E_{\text{th}}$. We observe that the proportion of the unclassified S-haplotypes decreases, but the proportion of the self-compatible ones increases with $E_{\text{th}}$. The figure is based on 16 independent runs for each $k$ value, with 1200 data points taken from each run, with a 25-generation interval between consecutive time points used for the analysis. All these data points were taken after the entire population descends from a single common ancestor and the amino acid frequencies have already reached their steady-state values. The total runtime for each is $2 \times 10^5$ generations. Parameter values: interaction energy thresholds $E_{\text{th}} = -6$. The remaining parameters are as in Table 2 main text.

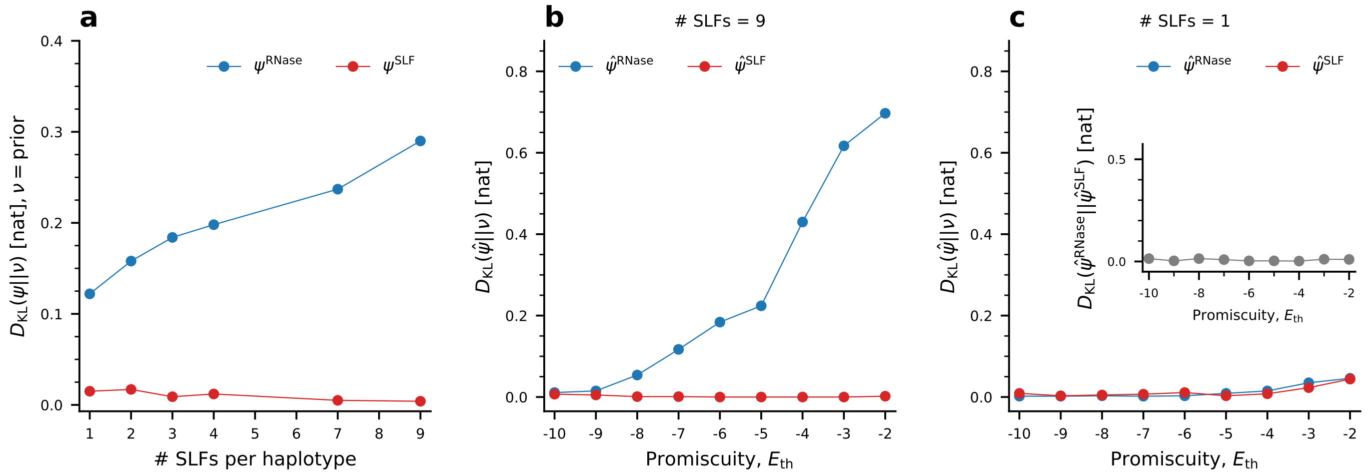


Figure S12: **The divergence of the RNase AA frequencies reduced relative to the prior increase with the number of SLF genes per S-haplotype and witn $E_{\text{th}}$ in both the full and the reduced models.** **(a)** The divergence of the the RNase AA frequencies relative to the prior increases with the number of SLF genes per S-haplotype. We illustrate the Kullback-Leibler divergence between the marginal AA distribution of evolved sequences $\psi$ and the prior AA distribution $\nu$ for RNases (blue) and functional SLFs (red) as a function of the number of SLFs per S-haplotype with fixed $E_{\text{th}}$ =-6. The calculations are based on 25 independent runs, using the last 1000 data points from each run with 25 generation intervals between consecutive time points (Methods). The total run time is 200000 generations. **(b)** The divergence of the reduced model including only selection against self-compatibility $\hat{\psi}$, relative to the prior $\nu$ against $E_{\text{th}}$. The divergence of the RNase frequencies (blue) increases with $E_{\text{th}}$, but the divergence of the SLF frequencies (red) is nearly zero. These results are in agreement with the perfect match between the full and reduced models, as shown in Fig. 4c. The calculations are based on 25 independent runs, using the last 600 data points from each run with 25 generation intervals between consecutive time points (Methods). The total run time is 150000 generations. **(c)** In the reduced model with only one SLF per S-haplotype, we obtain full symmetry between the RNase and SLF marginal distributions (compare to (b)). The inset shows the divergence between the RNase and SLF marginal distributions for the reduced model, again demonstrating the full symmetry in their behaviors. The calculations are based on 200 independent runs, using the last 400 data points from each run with 125 generation intervals between consecutive time points (Methods). The total run time is 200000 generations. Parameter values are as is Table 2 except for the numbers of SLFs per S-haplotype in (a) and (c).

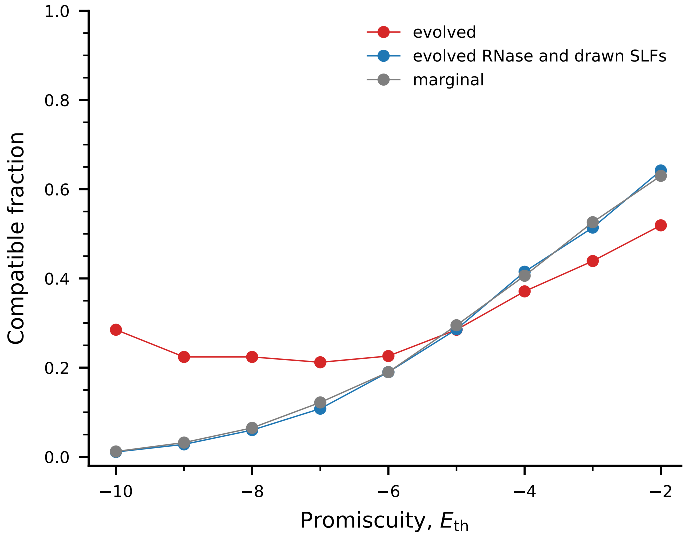


Figure S13: **Compatible fraction amongst RNase-SLF pairs in different scenarios**. We show here the compatible fraction among randomly paired RNase-SLF sequences evolved in simulation (red, 'evolved'). We show for comparison the compatible fraction between RNases evolved in the simulation and random SLF sequences drawn from the marginal distribution of evolved SLFs (blue), and the compatible fraction among random RNase and SLF sequences both drawn from their respective marginal distributions of the evolved sequences ('marginal', gray). The 'evolved' and 'marginal' curves are the same as in Fig. 4a. Both the blue and the gray curves closely match the red one for intermediate-to-high $E_{\text{th}}$, but deviate from it for lower $E_{\text{th}}$. This shows that for intermediate-to-high $E_{\text{th}}$ the sequences are mostly afected by biases in the single AA frequencies. In contrast, for low $E_{\text{th}}$ there is a significant contribution to local AA matches, that is not captured by the single AA frequencies, but only reflected in the joint RNase-SLF distribution. Drawing sequences from the marginals breaks these local matches. Note that the the blue and the gray curves nearly overlap. The reason is that it is sufficient to draw just one of the sequences (here the SLF) from the marginal to break these RNase-SLF dependencies. The figure is based on 16 independent runs with 1200 data points taken from each run, with a 25-generation interval between consecutive time points used for the analysis. All these data points were taken after the entire population descends from a single common ancestor and the amino acid frequencies have already reached their steady-state values. The total runtime for each is $2 \times 10^5$ generations. Parameter values: interaction energy thresholds $E_{\text{th}} = -6$. The remaining parameters are as in Table 2 main text.

### References

1. Vieira J, Rocha S, Vázquez N, López-Fernández H, Fdez-Riverola F, Reboiro-Jato M and Vieira CP. Predicting specificities under the non-self gametophytic self-incompatibility recognition model. *Front. Plant Sci.* **10**, 879 (2019).
